# ConnectoFM: A Foundation Model for Learning the Language of the Connectome

**DOI:** 10.64898/2026.06.06.730367

**Authors:** Abrar Rahman Abir, Anik Saha, Ruwad Naswan, Md. Shamsuzzoha Bayzid

**Affiliations:** Bangladesh University of Engineering and Technology

## Abstract

Accurate reconstruction of neural circuits from electron microscopy (EM) data is central to connectomics, yet modern datasets are now so large and heterogeneous that manual annotation and dataset-specific model retraining have become major challenges. While recent EM foundation models provide general visual representations, they are not specifically tailored to the connectomics domain, where preserving fine membrane boundaries and synaptic structures is essential to mitigate topological and connectivity errors. Here, we present ConnectoFM, the first foundation model for connectomics, pretrained on a diverse corpus of 1.7 million unlabeled EM images drawn from six species and 25 subdomains. ConnectoFM combines masked image modeling with contrastive alignment to learn robust visual representations directly from large-scale connectomics data. These representations organize EM images into biologically meaningful clusters across species, brain regions, developmental cohorts, and acquisition domains. Using frozen pretrained features with lightweight decoder heads, we transfer ConnectoFM to three important downstream tasks: binary segmentation, multiclass cell typing, and instance segmentation. Across 29 diverse datasets, including established benchmarks, ConnectoFM consistently outperforms existing EM foundation models and state-of-the-art methods that require task-specific training from scratch. With only 10% labeled data, ConnectoFM surpasses the baselines trained on 100% annotation budget, showing the superiority of ConnectoFM in low-data regimes. Improvements of ConnectoFM are especially pronounced for challenging and biologically important targets, including membranes, mitochondria, vesicles, post-synaptic densities and synapses, and remain strong in low-label settings. Extension to 3D volumetric segmentation and qualitative comparisons further show that ConnectoFM enables more accurate and biologically faithful performance across downstream tasks. These results establish ConnectoFM as a generalizable and data-efficient foundation model for connectomics and provide a scalable route towards more reliable neural circuit reconstruction.

## 1 Introduction

Connectomics seeks to map the structural wiring of the nervous system by reconstructing neural circuits at synaptic resolution, with the long-term goal of understanding how brain structure gives rise to function and computation. Advances in volume electron microscopy (EM) have made it possible to image neural tissue at nanometer resolution across increasingly large volumes, enabling reconstructions that range from canonical small nervous systems to modern large-scale brain datasets. Recent connectomic reconstruction efforts such as H01 in human cortex [1], MICrONS in mouse visual cortex [2], FlyWire in the adult *Drosophila* brain [3], and further studies spanning many other species [4–12] illustrate how rapidly connectomics is expanding in both scale and scope. This progress is scientifically transformative, but it also creates a major computational bottleneck: modern EM datasets routinely reach terabyte-to-petabyte scale, and manual analysis and proofreading no longer scale with data generation [13]. As a result, connectomics increasingly depends on computational methods that can transform raw EM images into reliable structural representations.

A central step in this pipeline is segmentation of neurons, cell membranes, synapses, and organelles, which enables downstream reconstruction and connectivity analysis [13]. Accurate segmentation remains difficult due to dense and heterogeneous ultrastructure, imaging artifacts, and the prevalence of merge and split errors in automated reconstructions [14]. Manual labeling and exhaustive proofreading are prohibitively time-consuming at the scale of modern EM volumes [13], and even small segmentation errors such as merges, splits, or boundary leaks can cascade into incorrect object topology and connectivity estimates. Moreover, segmentation models often require extensive labeled data and careful retraining to achieve high accuracy on each new volume, further amplifying the annotation burden [15]. Together, these challenges motivate computational approaches that reduce reliance on dense supervision and improve segmentation robustness and accuracy for large-scale connectomics.

Machine learning has become the dominant paradigm for connectomics segmentation, enabling automated delineation of neurons, membranes, synapses, and organelles from large-scale EM volumes. Prior work has extensively explored CNN based architectures, particularly U Net style models in both 2D and 3D settings for organelle and neuron segmentation [15–18]. Alternative formulations include affinity based prediction and agglomeration pipelines, often augmented with geometric supervision or graph based refinement to improve instance consistency at scale [19–21]. Iterative and instance oriented approaches such as flood filling and Mask R CNN style segmentation have also been investigated for high precision reconstruction and multi organelle settings [22, 23]. Despite strong progress, these methods still struggle with the scale and variability of EM volumes, and remain sensitive to ambiguous boundaries, anisotropy, and imaging artifacts, where residual merge and split errors can accumulate and require substantial downstream correction. Moreover, existing methods are typically engineered and trained for a narrow target setting, such as a specific organelle, tissue, imaging protocol, or annotation style, and often require substantial labeled data and careful retraining to reach high precision. Foundation models offer a more scalable alternative by learning general visual representations from large and diverse corpora, which can then be adapted to many downstream tasks with comparatively little additional supervision – for example, through lightweight fine-tuning or prompt-driven inference. This paradigm has shown strong success in different domains like medical imaging and natural images. Some prominent examples of foundation models for images include MedSAM [24], VISTA3D [25], MRI-CORE [26], ShapeMamba-EM [27], DendriteSAM [28], TriSAM [29] etc. They suggest that learning generalizable representations from large unlabeled or weakly labeled corpora can improve robustness and reduce annotation burden.

However, foundation models for 2D electron microscopy remain relatively underexplored and have only recently begun to emerge. This gap is notable because EM images present distinct challenges, including nanoscale ultrastructure, strong texture dominance, and substantial variability across preparation protocols and datasets, under which models trained for a single benchmark often fail to generalize. RETINA [30] takes an important step toward EM-specific pretraining by leveraging large-scale unlabeled EM data (CEM500K) and combining convolutional layers for local detail with transformer layers for global context, yielding improved cellular structure segmentation across multiple public EM datasets. Complementarily, microSAM [31] adapts the Segment Anything paradigm to microscopy by fine-tuning generalist promptable segmentation models for both light and electron microscopy, improving segmentation quality across diverse imaging conditions while enabling interactive and automatic annotation through a practical napari-based workflow. Together, these efforts suggest that EM foundation models can reduce dependence on dataset-specific training and provide more robust segmentation capabilities, motivating foundation model based approaches for connectomics EM images.

Despite the progress of foundation models in biomedical imaging and EM, a dedicated foundation model for connectomics EM images is still missing. This gap is critical because connectomics operates in a uniquely stringent regime, where segmentation must preserve fine membrane boundaries and synaptic structures over massive volumes, and small local errors can propagate into large topological and connectivity mistakes. A connectomics-oriented foundation model could therefore provide a shared, reusable representation that generalizes across species and brain regions, reduces reliance on dense supervision, improves robustness to artifacts and ultrastructural variability, and supports diverse downstream tasks.

To address this gap, we introduce ConnectoFM, a **F**oundation **M**odel for **Connecto**mics EM images that learns generalizable representations from large-scale unlabeled data and transfers effectively to downstream tasks. ConnectoFM is pretrained with a Vision Transformer based masked autoencoder that reconstructs masked image patches. In addition, we incorporate a contrastive alignment objective over augmented views of the same image, which promotes invariance to stochastic perturbations and stabilizes global representations. After pretraining, we transfer the frozen encoder to supervised tasks and train lightweight task-specific decoders on top of dense patch embeddings for pixel-level prediction.

A key design choice in ConnectoFM was whether to pretrain in 2D or 3D. Although 3D models can leverage inter-slice or axial context, they are often much more computationally expensive at connectomics scale, especially on anisotropic EM data, while adjacent serial EM slices are strongly correlated [32–35]. By contrast, 2D pretraining aligns naturally with mature vision architectures and highly scalable training pipelines. This choice is also well matched to practical connectomics workflows, where 3D segmentation is often built from strong 2D predictions followed by slice-to-slice registration, stacking, association, and agglomeration into volumetric reconstructions [32, 36–39]. Therefore, robust 2D representations provide a strong foundation for downstream 3D reconstruction and segmentation. For these reasons, we pretrain ConnectoFM on a large and diverse corpus of 1.7 million 2D EM images. However, we additionally evaluate ConnectoFM on 3D segmentation using a 2D-to-3D reconstruction pipeline with u-Segment3D [39]. Remarkably, although ConnectoFM is pretrained purely on 2D images, it remains effective when applied to 3D volumetric segmentation and outperforms general EM foundation models across benchmarks in 3D.

Our study makes the following key contributions. First, we introduce ConnectoFM, the first known foundation model designed specifically for connectomics, pretrained on a large and diverse corpus of 1.7 million unlabeled EM images curated from six species. Second, we show that ConnectoFM learns robust and biologically meaningful representations that organize connectomics data across multiple axes of variation, including species, brain regions, and developmental or life-stage cohorts. Third, we demonstrate the practical utility of these representations by adapting ConnectoFM to three important downstream tasks in connectomics such as binary segmentation, cell typing, and instance segmentation and evaluating it across 29 datasets, including well-established connectomics benchmarks. We find that ConnectoFM consistently and significantly outperforms state-of-the-art methods in binary segmentation and cell typing, while maintaining competitive accuracy in instance segmentation, reflecting its value for real-world connectomics workflows. Importantly, we show that ConnectoFM is substantially more data-efficient than other baselines in low-label regimes, enabling stronger downstream performance with reduced annotation requirements. Moreover, we analyze the representation geometry to explain the efficiency of ConnectoFM in low data regime. Fourth, ConnectoFM, despite being pretrained solely on 2D EM images, generalizes effectively to 3D volumetric segmentation via standard 2D-to-3D reconstruction pipelines, outperforming existing EM foundation models. Finally, we present case studies demonstrating the utility of our model and highlighting its advantages over existing EM foundation models and baseline methods.

## 2 Results

### 2.1 Overview of ConnectoFM

We developed ConnectoFM as a foundation model designed to learn transferable representations directly from large-scale unlabeled EM data in conectomics. The model is motivated by a central challenge in connectomics: although modern EM datasets span diverse species, brain regions, developmental stages, and acquisition settings, most downstream analyses still rely on dataset-specific models that require substantial annotation and retraining [13, 14]. ConnectoFM addresses this limitation by learning a shared representation space from broad connectomics data diversity, with the goal of supporting multiple downstream tasks from a single pretrained encoder. Fig. 1 presents an overview of our study, encompassing the ConnectoFM training architecture, the diverse dataset sources, and the downstream adaptation pipeline.

**Figure 1:**
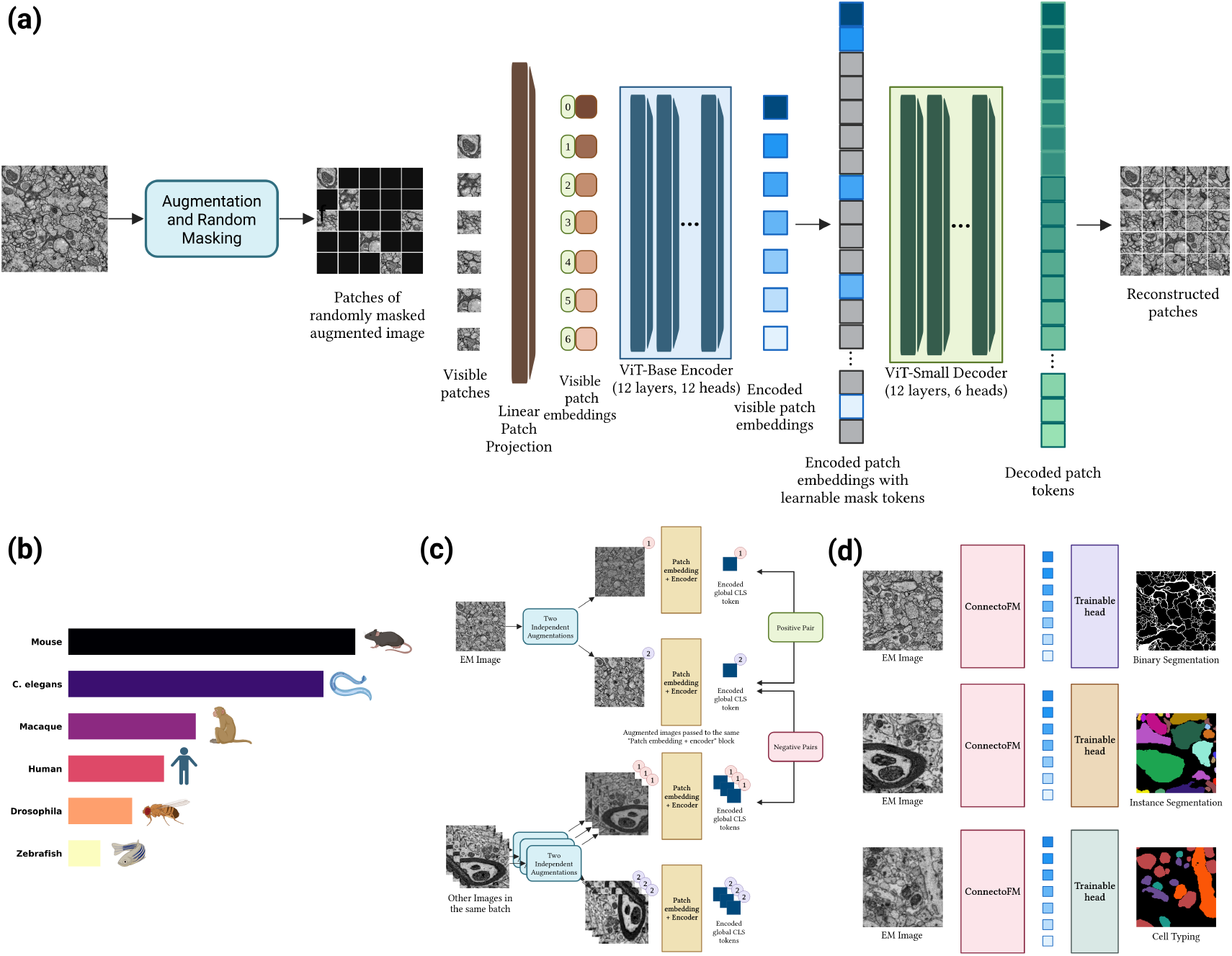
Overview of ConnectoFM. **a**, Pretraining architecture of ConnectoFM. An input EM image is augmented and randomly masked at the patch level. Visible patches are encoded, combined with learnable mask tokens, and passed to a decoder to reconstruct the masked content. **b**, Composition of the ConnectoFM pretraining corpus, comprising 1.7 million images from six diverse species. **c**, Contrastive alignment during pretraining. Two independently augmented views of the same EM image are processed by a shared patch embedding and encoder network, and their global representations are aligned as a positive pair. By contrast, views derived from different images form a negative pair. The contrastive loss pulls positive pairs closer and pushes negative pairs farther apart in latent space. **d**, Transfer of the pretrained encoder to downstream connectomics tasks using lightweight task-specific heads, including binary segmentation, instance segmentation, and cell typing. This figure was created in BioRender.com.

ConnectoFM is pretrained on 1.7 million unlabeled EM images collected across six species and 25 connectomics subdomains (see Section 5 (Materials and Methods) “Curation of the pretraining corpus”). To capture both local ultrastructural detail and broader contextual regularities, we train the model with a Vision Transformer-based masked autoencoding objective, which reconstructs masked image patches from visible context. We further augment this objective with contrastive alignment so that images with related biological or acquisition context are encouraged to map to nearby representations, while dissimilar images are separated in feature space. Together, these objectives enable ConnectoFM to learn embeddings that are not only reconstructive, but also discriminative and transferable across heterogeneous connectomics domains. Additional details of the model architecture and pretraining objectives are provided in Section 5 (Materials and Methods) “Pretraining of ConnectoFM”.

After pretraining, we transfer the frozen ConnectoFM encoder to downstream tasks using lightweight task-specific convolutional decoder heads. This design allows us to test whether the pretrained representation itself contains general information useful for connectomics, without relying on extensive end-to-end retraining. We evaluate ConnectoFM on three representative downstream settings: binary segmentation, multiclass cell typing, and instance segmentation. These tasks were chosen to span distinct prediction regimes, from dense boundary-sensitive segmentation to higher-level semantic discrimination and object-level partitioning. The adaptation protocols across the three downstream tasks and the benchmark datasets are further described in the Materials and Methods section “Downstream adaptation” and “Datasets used for downstream evaluation”. Detailed definitions of the evaluation metrics, together with the average per-sample inference times for these downstream tasks, are provided in the Supplementary Material (Section 1 and Table S4).

Using this framework, we examine three questions (discussed in subsequent sections). First, we ask whether large-scale pretraining on connectomics-specific data yields biologically meaningful representation structure across species and domains. Second, we test whether these learned features improve downstream performance relative to strong supervised and foundation-model baselines. Third, we assess whether the benefits of ConnectoFM persist in practically important regimes, including limited-label training and 3D volumetric inference.

### 2.2 ConnectoFM Learns Meaningful Representations Across Diverse Domains

To assess the quality of the representations learned during pretraining, we examine the domain organization and reconstruction capability of ConnectoFM on the held out portion (4%) of the pretraining corpus. We first analyze the geometric structure of ConnectoFM encoder embeddings using two complementary views: (i) a joint projection that includes all species and their subdomains (Fig. 2a-b), and (ii) species conditioned projections that isolate within species variation (Fig. 2c). Here, *domains* correspond to species (C. elegans, Drosophila, Human, Macaque, Mouse, Zebrafish), and *subdomains* capture biologically and experimentally meaningful partitions within each species, including developmental stages (L1 newborn, L1 early hours, late L1 stage, mid L2 stage, L2 stage, L3 stage, young adult, and adult for C. elegans), anatomical regions (visual cortex, CA1 hippocampus, cerebellum for Mouse and Cortex for Human), and layer specific samples at different time points (layer 2/3 day 105, layer 2/3 day 523, layer 4 day 6, layer 4 day 105, layer 4 day 523 for Mouse and layer 2/3 day 7, layer 2/3 day 75, layer 2/3 day 3000, and layer 4 day 3000 for Macaque). We further quantify local neighbourhood consistency with a kNN evaluation on the same embedding space, where higher accuracy indicates that samples sharing the same subdomain label are concentrated in local neighbourhoods.

**Figure 2:**
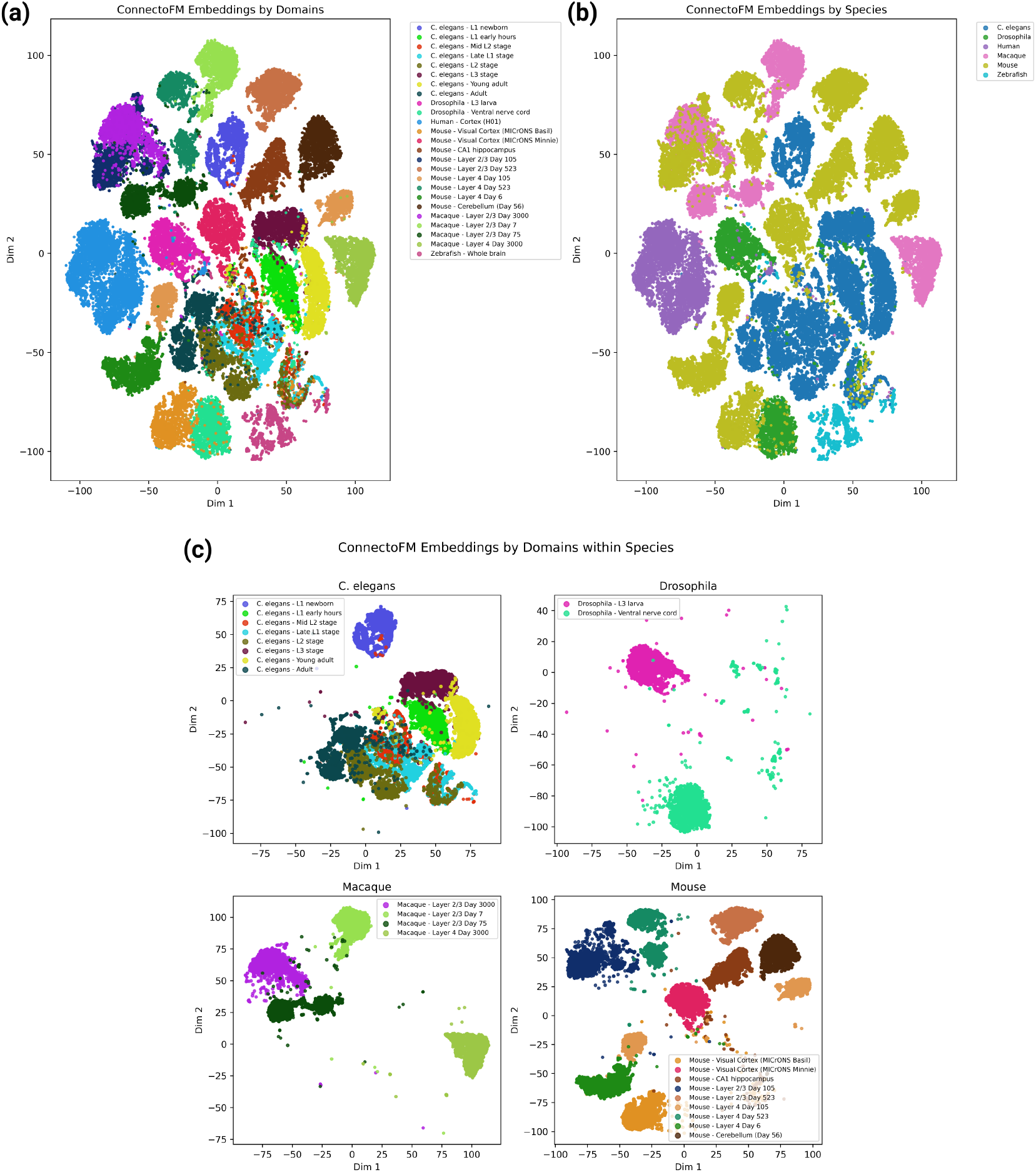
Embedding structure learned by ConnectoFM during pretraining across species and connectomics subdomains. **a**, Two-dimensional TSNE projection of embeddings for all samples pooled across species and subdomains, colored by subdomain. Subdomains include *C. elegans* developmental stages, *Drosophila* cohorts (L3 larva, ventral nerve cord), human cortex, mouse brain regions and cortical layers across timepoints, macaque cortical layers across timepoints, and zebrafish whole brain. **b**, The same pooled embedding space, colored by species, showing species-level organization across the pretrained representation. **c**, Species-specific projections illustrating within-species organization of subdomains, where clusters align with developmental stage, brain region, cortical layer, and acquisition cohort, reflecting structured representations learned from the pretraining corpus.

#### Joint embedding structure across species and subdomains

In the joint projection (Fig. 2a), ConnectoFM organizes samples into coherent clusters that align with the full set of connectomics domains and subdomains: C. elegans developmental stages (L1 newborn, L1 early hours, late L1 stage, mid L2 stage, L2 stage, L3 stage, young adult, adult), Drosophila (L3 larva, ventral nerve cord), Human (cortex), Mouse (visual cortex MICrONS Basil, visual cortex MICrONS Minnie, CA1 hippocampus, layer 2/3 day 105, layer 2/3 day 523, layer 4 day 6, layer 4 day 105, layer 4 day 523, cerebellum day 56), Macaque (layer 2/3 day 7, layer 2/3 day 75, layer 2/3 day 3000, layer 4 day 3000), and Zebrafish (whole brain). Remarkably, the corresponding kNN accuracy over the joint label space is 0.966 (Table 1), indicating that embeddings preserve strong neighbourhood level separability even when all species and subdomains are mixed together. In the species colored view (Fig. 2b), embeddings exhibit clear species level macro structure, with each species forming compact islands while still retaining internal fine scale variation, suggesting that the representation captures both global distribution shifts across species and within species biological structure.

**Table 1:** kNN accuracy of ConnectoFM embeddings across joint and species-specific label spaces. Higher values indicate stronger local neighbourhood consistency, such that samples sharing the same subdomain label are concentrated in nearby regions of the embedding space.

| Setting | Label space | kNN accuracy |
| --- | --- | --- |
| Joint embedding | All species and subdomains | 0.966 |
| Species-specific | *Drosophila* subdomains | 0.996 |
| Species-specific | *Macaca mulatta* subdomains | 0.997 |
| Species-specific | *Mus musculus* subdomains | 0.995 |
| Species-specific | *C. elegans* developmental subdomains | 0.945 |

#### Within species subdomain organization

Fig. 2c probes whether ConnectoFM retains meaningful structure *within* a species rather than relying only on coarse species separation. We further report species-specific kNN accuracies in Table 1. For Drosophila, the two subdomains (L3 larva and ventral nerve cord) form two well separated clusters, reflected by a very high kNN accuracy of 0.996. For Macaque, subdomains defined by cortical layer and timepoint (layer 2/3 at day 7, day 75, day 3000, and layer 4 at day 3000) clearly separate into distinct groups, with layer 4 day 3000 forming a clearly isolated cluster relative to layer 2/3 samples, and the kNN accuracy reaches 0.997, providing near-perfect quantitative evidence of subdomain separability. For Mouse, embeddings clearly separate multiple anatomical regions and cohorts, including MICrONS visual cortex (Basil and Minnie), CA1 hippocampus, cerebellum day 56, and layer specific samples (layer 2/3 at days 105 and 523; layer 4 at days 6, 105, and 523), with a kNN accuracy of 0.995, further confirming strong separability. In contrast, C. elegans shows more overlap among developmental subdomains (L1 newborn, L1 early hours, late L1 stage, mid L2 stage, L2 stage, L3 stage, young adult, adult), consistent with a smoother progression across stages, and correspondingly exhibits the lowest within species kNN accuracy at 0.945. Taken together, the projections and kNN scores indicate that ConnectoFM learns embeddings that are strongly structured across species and remain discriminative for fine grained connectomics subdomains such as brain regions, cortical layers, developmental stages, and dataset cohorts.

#### Accurate reconstruction of augmented EM images from heavily masked input

To assess whether the pretraining objective was achieved, we next examine ConnectoFM’s ability to reconstruct augmented EM images from inputs in which 75% of patches were masked (Fig. 3). Each image is first transformed by the same augmentation pipeline used during pretraining, including resizing, random resized cropping, and optional horizontal flipping. The model must then infer the missing content from only sparse visible patches. Remarkably, across all six domains, ConnectoFM reconstructs the perturbed views with high fidelity recovering delicate contours and small structural features that are only weakly visible in the masked input. This capability is clear in Fig. 3 when the reconstructed images (model predicitons) are compared against their corresponding pre-masking versions (reconstruction targets), where the model’s ability to faithfully recover the underlying structures is evident. This suggests that ConnectoFM learns representations that extend beyond coarse image statistics to encode biologically meaningful ultrastructural features, thereby providing a strong basis for downstream connectomics tasks.

**Figure 3:**
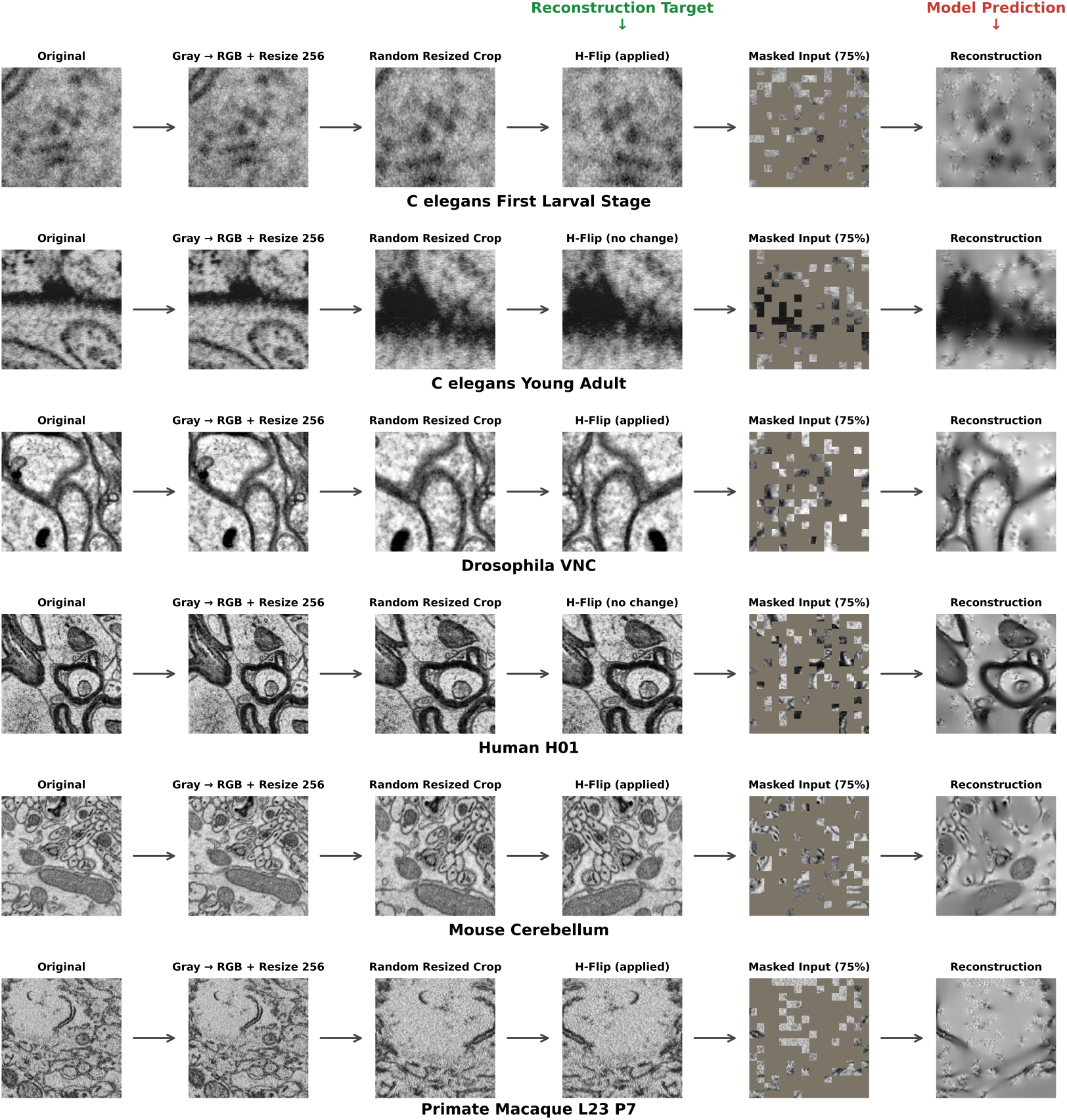
Reconstruction of augmented EM views from heavily masked inputs by ConnectoFM. Each row shows one representative example from a different connectomics domain: *C. elegans* first larval stage, *C. elegans* young adult, *Drosophila* ventral nerve cord (VNC), human H01 cortex, mouse cerebellum, and primate macaque layer 2/3 cortex. For each example, we show the original image, grayscale-to-RGB conversion with resizing to 256 *×* 256, a random resized crop, the optionally flipped augmented EM view used as the pretraining target, the corresponding masked input with 75% of patches removed, and the reconstruction predicted by ConnectoFM. The reconstruction objective is therefore to recover the augmented target view (fourth column). Across diverse species and imaging domains, the model recovers both the global morphology and fine ultrastructural patterns of the target view, suggesting that pretraining captures representations that retain local detail while preserving broader spatial context.

### 2.3 ConnectoFM Performs Accurate Binary Segmentation Across Organelles and Datasets

We evaluate ConnectoFM on binary segmentation across 22 connectomics EM datasets spanning diverse organelles including mitochondria, membranes, synapses, vesicles, post-synaptic densities (PSD), axons, and mitochondrial boundaries. Supplementary Table 2 provides details on the datasets used in binary segmentation. We compare ConnectoFM against three representative baselines: RETINA [30] and microSAM[31], which are foundation model based methods developed for general EM imagery, and Pytorch Connectomics UNet (PyTC UNet) [40], a UNet-based baseline widely used in connectomics community [41]. For ConnectoFM and RETINA, we freeze the corresponding encoder weights and train the decoder in a supervised manner. For PyTC UNet, we train the model from scratch on each dataset. For microSAM, we adopt a point-prompt-based evaluation protocol consistent with prior studies [31, 42–44]. Specifically, for each 512 *×* 512 patch of an image, we randomly sample 10 positive prompt points from ground-truth foreground pixels and 10 negative prompt points from ground-truth background pixels, and provide these prompts to microSAM during inference. To account for stochastic variation, we repeat this sampling procedure across 10 different seeds and report the mean performance. Fig. 4a compares Dice scores across all datasets for the four methods. Comparisons on all datasets using various metrics, including precision, recall, and IoU, are provided in the Supplementary Material (see Figures S1-S4). In this section, we focus our discussion on Dice scores, because Dice is particularly well-suited for binary segmentation with class imbalance, directly measuring overlap between predicted and ground truth masks.

**Figure 4:**
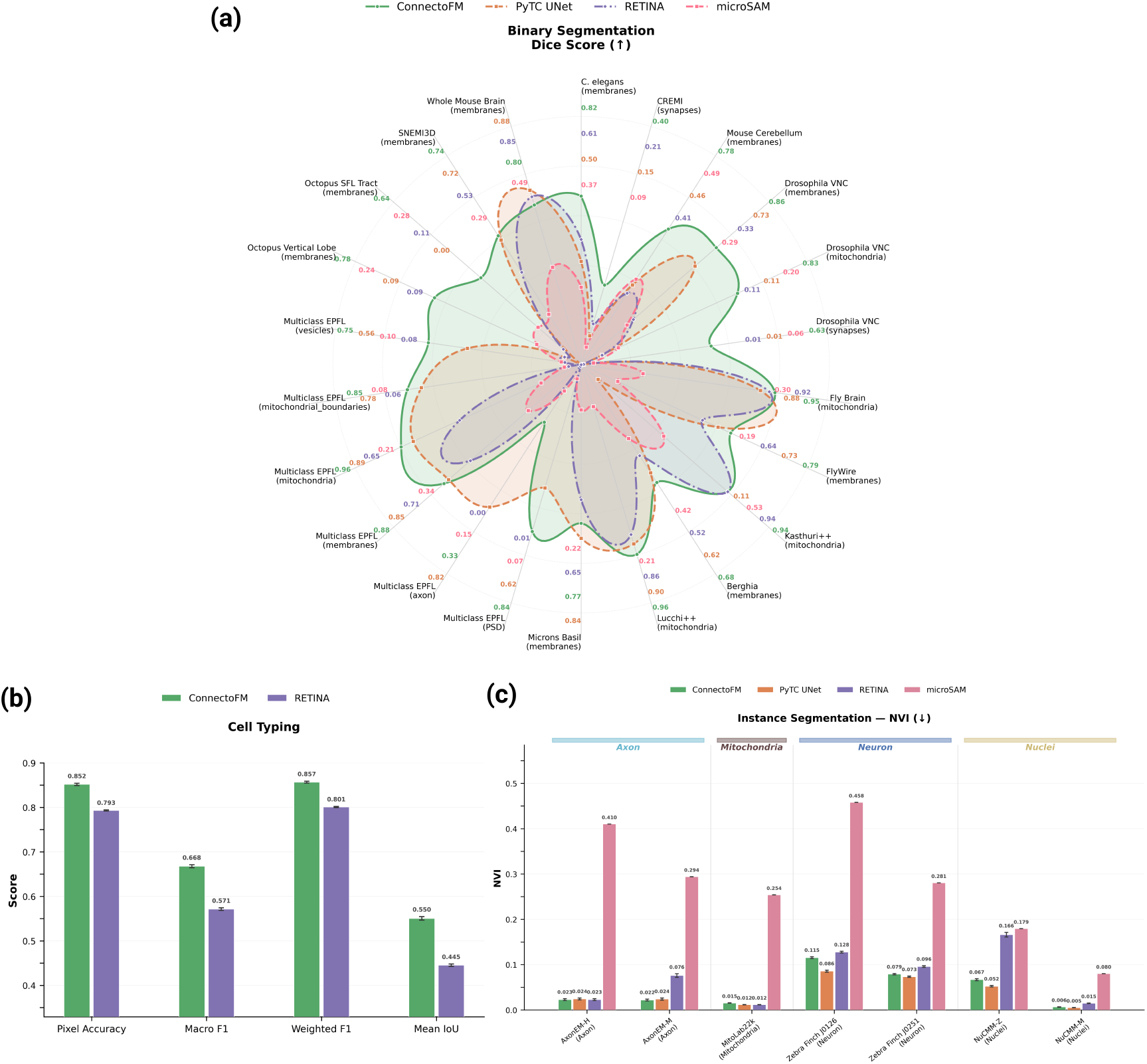
Comparison of ConnectoFM with baseline methods across three downstream tasks. **a**, Dice-score comparison of ConnectoFM, PyTC UNet, RETINA, and microSAM on binary segmentation across 22 datasets spanning diverse species and organelles. Dice scores are shown in ascending order radially outward. **b**, Cell-typing performance of ConnectoFM and RETINA on the Zebra Finch J0251 dataset, measured by pixel accuracy, macro F1, weighted F1, and mean IoU. Error bars indicate variability across five independent training runs with different seeds. For all metrics in **b**, higher values indicate better performance. **c**, Comparison of normalized variation of information (NVI) between ConnectoFM, PyTC UNet, RETINA, and microSAM on instance segmentation across 7 diverse datasets. Error bars indicate variability in per-sample metric values across the evaluation set. Lower values indicate better performance.

Averaged across all datasets and organelles, ConnectoFM achieves a mean Dice score of 0.771, substantially outperforming PyTC UNet (0.556), RETINA (0.423), and microSAM (0.256). This corresponds to an average Dice improvement of 38.67% over PyTC UNet, 82.53% over RETINA, and 201.40% over microSAM. Notably, ConnectoFM attains the highest Dice score on 20 of the 22 dataset–task pairs, consistently surpassing the competing methods. All improvements are statistically significant (*p <* 0.01) with the exception of Mouse cerebellum (membranes) against PyTC UNet (*p* = 0.029) and RETINA (*p* = 0.024), Berghia (membranes) against PyTC UNet (*p* = 0.129) and RETINA (*p* = 0.251), and Multiclass EPFL (axons) against PyTC UNet (*p* = 0.192). Importantly, these gains remain consistent across datasets with diverse imaging conditions and varying anatomical complexity, indicating that ConnectoFM generalizes well beyond any single dataset or organelle type.

On membrane segmentation across 11 diverse datasets, ConnectoFM attains an average Dice score of 0.78, markedly outperforming PyTC UNet (0.58), RETINA (0.50), and microSAM (0.33). This clear margin highlights its ability to construct continuous, non-porous membrane masks, suggesting that the ConnectoFM encoder captures both local detail and broader contextual structure effectively.

The advantage of ConnectoFM becomes even more pronounced in mitochondria segmentation. Averaged across five diverse datasets, including established benchmarks such as Lucchi++, Kasthuri++, and the Fly Brain Mitochondria benchmark, ConnectoFM achieves a Dice score of 0.93, substantially surpassing PyTC UNet (0.58), RETINA (0.69), and microSAM (0.29). This strong margin indicates that ConnectoFM captures both mitochondrial shape and internal consistency more reliably than the baseline methods, enabling accurate reconstruction of abundant, spatially distributed, and morphologically diverse organelles in EM images. On the related task of segmenting mitochondrial boundaries in the Multiclass EPFL dataset, ConnectoFM further attains a Dice score of 0.85. By comparison, PyTC UNet, RETINA, and microSAM achieve Dice scores of 0.78, 0.06, and 0.08, respectively. These results indicate that the learned representations are sensitive not only to organelle interiors but also to fine structural boundaries.

Accurate segmentation of synapses, axons, and post-synaptic densities (PSDs) is both challenging and essential in connectomics, as these structures are fundamental to the faithful reconstruction of brain connectivity networks [45–47]. Among them, synaptic structures are particularly difficult to segment because they are extremely sparse and exhibit substantial morphological variability [14, 48]. Despite these challenges, ConnectoFM achieves average Dice scores of 0.84 for PSD segmentation and 0.52 for synapse segmentation, underscoring its ability to generalize to small and spatially sparse structures. By contrast, RETINA attains Dice scores of only 0.01 for PSDs and 0.11 for synapses, while microSAM reaches only 0.07 for PSDs and 0.07 for synapses. Synapse segmentation is especially challenging, with all baseline methods performing poorly overall, reaching average Dice scores of 0.11 for RETINA, 0.08 for PyTC UNet, and 0.07 for microSAM.

On vesicle segmentation, ConnectoFM achieves a Dice score of 0.75, outperforming PyTC UNet (0.56), RETINA (0.08), and microSAM (0.10). This result further highlights the difficulty that baseline methods face in segmenting small, densely packed structures. Axon segmentation is the only notable exception to this overall trend. On this task, PyTC UNet achieves the best Dice score (0.82), while ConnectoFM (0.33) outperforms microSAM (0.15) and substantially exceeds RETINA, which fails entirely on this task (0.00). These results indicate that the advantage of ConnectoFM is strong but not uniform across all structure types.

Thus, ConnectoFM delivers consistent and often substantial gains across binary segmentation tasks, outperforming both a UNet trained from scratch and existing foundation-model-based baselines. Notably, although RETINA and microSAM are also trained on EM images, their inferior segmentation performance on the same modality further underscores the importance of connectomics-specific representation learning.

### 2.4 ConnectoFM Enables Reliable Multiclass Cell-Type Classification from EM Images

Dense cell-type classification of neuronal and non-neuronal cells from EM images is another critical downstream task for connectomics because a connectome becomes biologically interpretable only when reconstructed nodes are assigned molecularly and morphologically meaningful identities, enabling cell-type-resolved connectivity matrices, circuit motif analysis, and mechanistic models of computation [49–52]. Therefore, we next evaluate ConnectoFM on multiclass cell typing in the Zebra Finch J0251 dataset. The task requires predicting, for each pixel in the EM image, the cell-type label corresponding to the cell instance that pixel belongs to, yielding a dense multiclass semantic labelling map. The dataset contains 17 annotated classes, split into three non-neuronal types and fourteen neuronal types. We exclude two classes : FRAG and OLIGO, because they have zero support (proportion 0%) in the dataset. The neuronal labels capture prominent populations such as medium spiny neurons (MSN), dopaminergic neurons (DA), several interneuron subtypes (INT1, INT2, INT3), and basal-ganglia-associated groups (GPe, GPi, STN, TAN) whereas the non-neuronal labels correspond to astrocytes (ASTRO), microglia (MICRO), and oligodendrocytes (OLIGO). We compare ConnectoFM against another foundation model based method RETINA on multiclass cell-type classification, reporting both aggregate performance in Fig. 4b and detailed class-wise results across annotated cell types in Table 2. We evaluate performance using pixel accuracy, macro F1, weighted F1, and mean IoU.

**Table 2:**
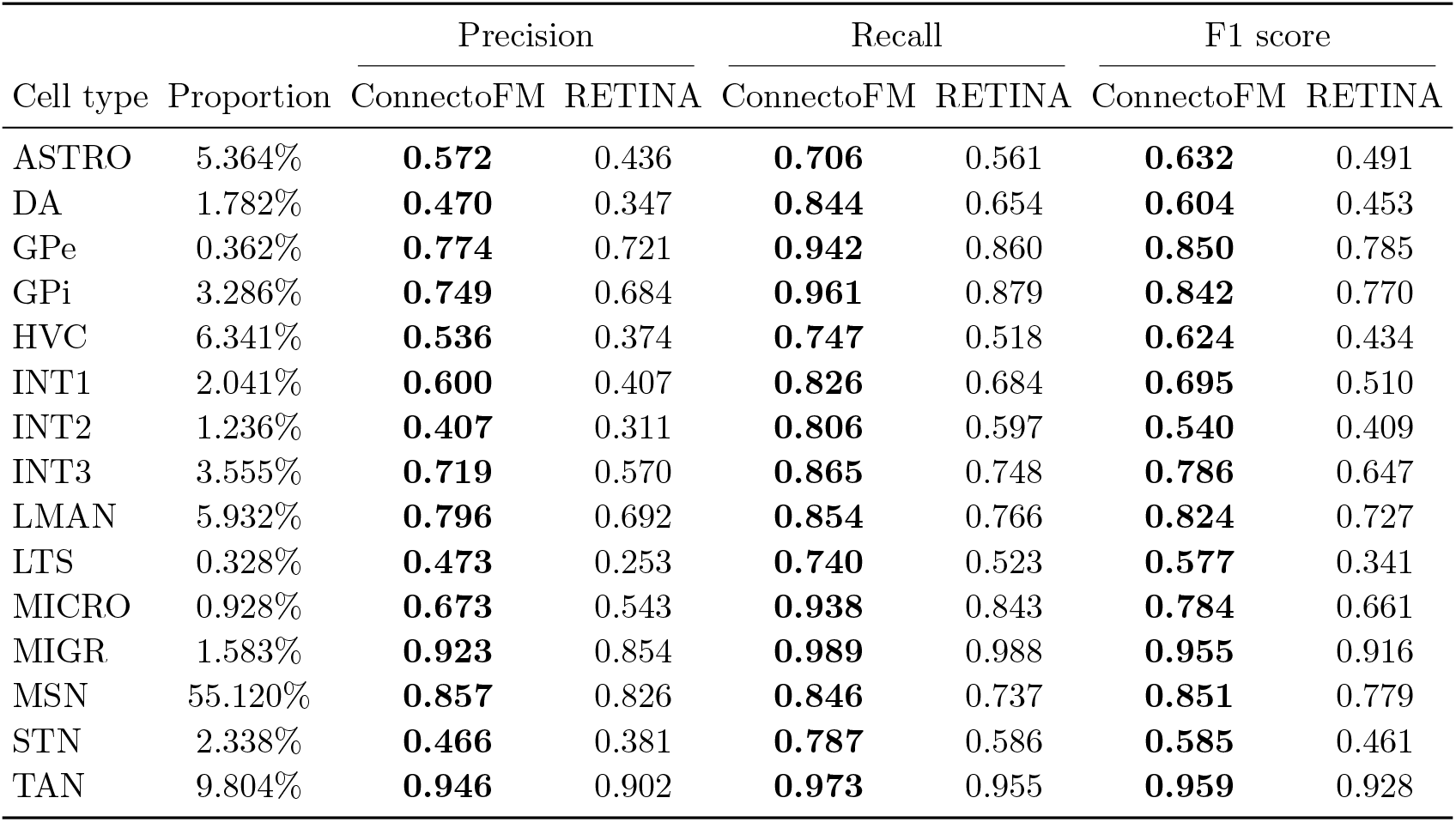
Cell-type-wise comparison of ConnectoFM and RETINA on cell type classification in the Zebra Finch J0251 dataset. FRAG and OLIGO are excluded because they have zero support in the dataset. The foreground proportion indicates the percentage of each cell type among all foreground pixels in the dataset. For each metric, the better score is shown in bold.

At the aggregate level, ConnectoFM consistently outperforms RETINA across all cell-typing metrics (Fig. 4b). ConnectoFM achieves a pixel accuracy of 0.852, compared with 0.793 for RETINA, indicating a substantially higher fraction of correctly classified pixels overall. The advantage is even more pronounced on class-balanced metrics. ConnectoFM attains a Macro F1 score of 0.668, whereas RETINA reaches 0.571, corresponding to a relative improvement of 16.9%. This gap points to markedly better handling of minority and low-support cell classes, while RETINA appears more biased toward dominant cell types. ConnectoFM also improves Weighted F1 from 0.801 to 0.857, showing that gains on underrepresented classes do not come at the expense of performance on abundant classes. A similar pattern is reflected in mean IoU, which increases from 0.445 for RETINA to 0.550 for ConnectoFM. Notably, all of these improvements are statistically significant (*p* ≪ 0.001).

Notably, ConnectoFM consistently and substantially outperforms RETINA on every evaluation metric for every cell type (see Table 2). For the most dominant neuronal type medium spiny neuron (MSN) with proportion of 55.120%, ConnectoFM consistently surpasses RETINA. In this case, ConnectoFM achieves an F1 score of 0.851, compared with 0.779 for RETINA. For the second most dominant cell type tonically active neuron (TAN) with proportion of 9.804%, ConnectoFM yields F1 score of 0.959, while RETINA attains 0.928. ConnectoFM further improves classification of other major neuronal groups, including GPe (0.850 versus 0.785), GPi (0.842 versus 0.770), LMAN (0.824 versus 0.727), and STN (0.585 versus 0.461). These results show that the learned representation remains robust even for large and heterogeneous neuronal populations.

The advantage becomes even clearer for more challenging subtypes. For the most challenging interneuron class INT2, ConnectoFM attains F1 score of 0.540, markedly higher than RETINA’s 0.409. Similar gains are observed for dopaminergic neurons (DA; 0.604 versus 0.453), HVC neurons (0.624 versus 0.434), and LTS cells (0.577 versus 0.341). This pattern suggests that ConnectoFM captures discriminative ultrastructural cues that are essential for resolving fine-grained neuronal identities.

A similar trend holds for non-neuronal and transitional cell states. Astrocyte classification improves from an F1 of 0.491 with RETINA to 0.632 with ConnectoFM, and microglia classification improves from 0.661 to 0.784. For migratory or immature cells (MIGR), ConnectoFM reaches 0.955, compared with 0.916 for RETINA. Taken together, these results indicate that ConnectoFM yields more reliable multiclass cell typing across both abundant and challenging cell populations.

Overall, these results show that ConnectoFM consistently outperforms RETINA in both aggregate and class-wise evaluation. The gains extend well beyond overall pixel accuracy, with particularly strong improvements in Macro F1, indicating more reliable recognition of rare and fine-grained cell types. By preserving strong performance on dominant classes while markedly improving minority-class classification, ConnectoFM delivers a more balanced and biologically faithful framework for dense EM-based cell typing.

### 2.5 ConnectoFM Achieves Strong Instance Segmentation Performance

We next evaluate ConnectoFM on instance segmentation, which requires separating individual object instances rather than only predicting foreground masks. This is a substantially more demanding setting because correct predictions must preserve object identity, prevent merges between adjacent instances, and avoid over-splitting single objects into multiple fragments. We conduct this evaluation across 7 datasets and 4 organelle categories. We compare ConnectoFM against three baselines: RETINA, microSAM, and PyTC UNet. Our main goal here is to determine whether a connectomics-specific pretrained representation translates into better instance-level understanding. We evaluate performance using Variation of Information split and merge terms (VOI-split and VOI-merge), their sum (VOI-total), Normalized Variation of Information (NVI), and Rand F-score. Higher Rand F-score indicates better instance agreement, whereas lower VOI and NVI indicate fewer topological and clustering-level errors. Figure 4c compares NVI scores across the methods, and the remaining metrics are reported in the Supplementary Material (see Figures S5-S9). In this section, we focus our discussion on NVI, as this metric is particularly well suited for instance segmentation by capturing discrepancies in both over-segmentation and under-segmentation through an information-theoretic measure of partition similarity. All performance differences observed in this task are statistically significant according to a paired *t*-test (*p <* 0.01).

Across all datasets and organelles, ConnectoFM achieves VOI-total = 0.7236, NVI = 0.0468, and Rand F-score = 0.8872. This places ConnectoFM clearly ahead of both foundation model baselines and second overall. In particular, compared to RETINA, ConnectoFM improves Rand F-score from 0.8322 to 0.8872, reduces VOI-total from 1.1311 to 0.7236, and reduces NVI from 0.0736 to 0.0468. This corresponds to a 36.3% reduction in NVI and a 36.0% reduction in VOI-total relative to RETINA. The margin over microSAM is even larger: NVI is reduced from 0.2796 to 0.0468, a 83.2% reduction, while VOI-total drops from 2.5202 to 0.7236, a 71.3% reduction. Rand F-score also increases substantially from 0.5985 to 0.8872. These aggregate results strongly support the advantage of ConnectoFM over the two FM baselines. This is particularly meaningful because RETINA and microSAM are pretrained on general EM imagery, whereas ConnectoFM is specialized for connectomics. The large gap therefore suggests that domain specificity matters strongly for instance segmentation: representations tuned to generic EM content are not sufficient to model the dense packing, fine boundaries, and highly entangled morphologies that are characteristic of connectomics volumes.

At the same time, ConnectoFM remains highly competitive with PyTC UNet. PyTC achieves the strongest overall results, with NVI = 0.0396, but the gap to ConnectoFM remains relatively modest compared to the much larger separation from RETINA and microSAM. For example, the absolute NVI gap between PyTC and ConnectoFM is only 0.0073, whereas the gap between ConnectoFM and RETINA is 0.0267, and the gap between ConnectoFM and microSAM is 0.2327. Similarly, the Rand F-score difference between PyTC and ConnectoFM is only 0.0201. Thus, although PyTC ranks first overall, ConnectoFM remains very close and competitive, while being decisively stronger than the other two FM-based baselines.

The organelle-wise analysis provides a more fine-grained picture and further highlights the strengths of ConnectoFM. For Axon instance segmentation, ConnectoFM is the strongest method across all major metrics. It achieves NVI = 0.0224. Remarkably, this is better than PyTC UNet on each of these metrics, and substantially better than RETINA and microSAM. In particular, Axon NVI is reduced from 0.0494 for RETINA to 0.0224 for ConnectoFM, and from 0.3523 for microSAM to 0.0224, demonstrating a significant reduction in instance-level clustering errors. This result is important because axonal structures are elongated, densely packed, and prone to fragmentation and identity switches, making them especially challenging for instance segmentation.

For Neuron instance segmentation, ConnectoFM again shows a clear advantage over the FM baselines. It achieves NVI = 0.0972, compared to NVI = 0.1118 for RETINA, and NVI = 0.3693 for microSAM. Rand F-score also improves from 0.6877 for RETINA and 0.4035 for microSAM to 0.7259 for ConnectoFM. These margins indicate that ConnectoFM better preserves neuronal instance identity while reducing both merge and split errors. Although PyTC remains best on this organelle, ConnectoFM is again clearly separated from the FM baselines and remains reasonably close to the leading score. For Nuclei, the same trend holds.

For Mitochondria, both ConnectoFM and PyTC remains extremely accurate. ConnectoFM achieves NVI = 0.0153, where PyTC achieves NVI = 0.0118. On the other hand, while RETINA remains competitive, microSAM yields much lower performance in Mitochondria instance segmentation.

Overall, these results show that ConnectoFM provides the strongest instance segmentation performance among the foundation model based methods and does so by a substantial margin. While PyTC UNet achieves the best aggregate scores, ConnectoFM remains very close and competitive, and is in fact the best method on Axon. These findings indicate that connectomics-specific pretraining is highly beneficial for instance-level reasoning, yielding much stronger instance separation than general EM foundation models.

### 2.6 ConnectoFM Demonstrates Superiority in Low Data Regime

We performed an explicit low-data regime study for binary segmentation motivated by the practical challenges of annotation in connectomics. High-resolution EM volumes are extremely costly to label, requiring expert annotators and substantial time to delineate fine ultrastructural details such as membranes, synapses, and organelles. As a result, in many realistic settings, only a small fraction of the data can be densely annotated. This creates a need for models that remain effective under limited supervision. To evaluate this, we consider a setting where the encoder is kept frozen and pretrained, and only the segmentation head is trained using different fractions of the labeled training set. Specifically, for each dataset and organelle, we trained the segmentation head with the full training set and with reduced subsets containing 80%, 60%, 40%, 20%, and 10% of the original training data, while keeping the test protocol unchanged. This experiment isolates a central question: *how much labeled supervision is actually needed once the representation is already learned?* We compared ConnectoFM against two baselines under the same protocol: RETINA, a foundation model based method pretrained on general EM imagery, and PyTC UNet, a standard supervised U-Net based baseline. We present the results in Figure 5.

**Figure 5:**
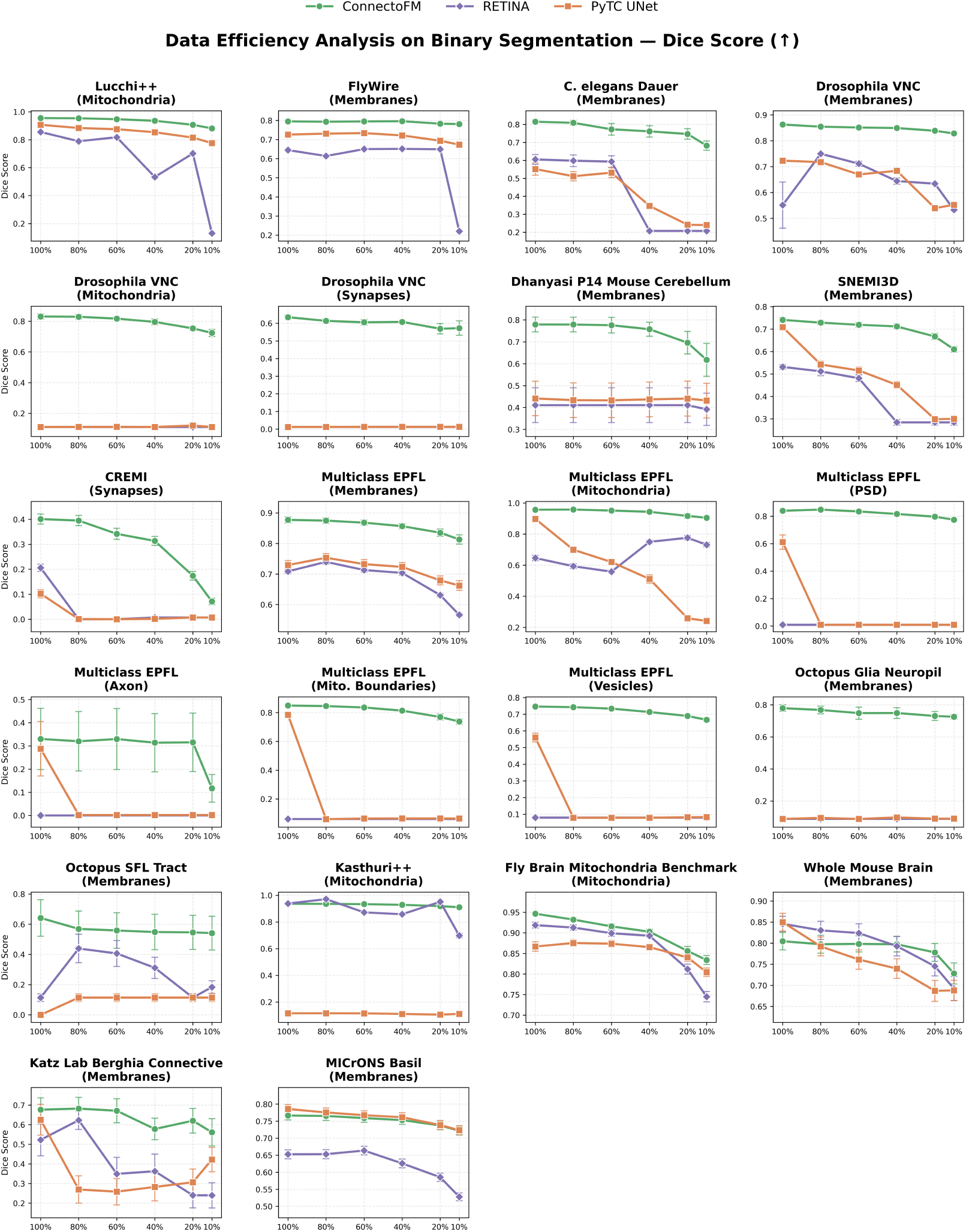
Data efficiency analysis of binary segmentation. For each task, only the segmentation head is trained while the encoder is kept frozen, using varying fractions of the labeled training data (100%, 80%, 60%, 40%, 20%, 10%). ConnectoFM consistently maintains high Dice scores even at extreme low-data settings (10–20%), showing only mild degradation relative to full-data training. In contrast, RETINA and PyTC UNet exhibit substantial performance drops as training data decreases, often failing on challenging organelles such as synapses, vesicles, and PSD. Notably, ConnectoFM trained with 10–20% data frequently matches or surpasses baseline performance trained with 100% data. Error bars indicate variability across test samples.

Our key finding is that ConnectoFM remains highly effective even in the extreme low data regime. The performance drop from full supervision to 20% and 10% labeled data is remarkably small, whereas both RETINA and PyTC UNet deteriorate much more severely. This behavior is consistent with the role of the ConnectoFM encoder: since it is pretrained specifically on connectomics EM and then frozen, the segmentation head is not forced to learn connectomics features from scratch. Instead, it only needs to learn a relatively simple task specific readout from already rich and task aligned representations. In contrast, RETINA, although foundation model based, is not specialized for connectomics and must adapt more generic EM features to dense connectomic structures using limited labels. PyTC UNet is even more label hungry because it must learn both feature extraction and task mapping directly from the segmentation supervision. As a result, ConnectoFM has much lower sample complexity.

This effect is already clear from the average Dice across all evaluated dataset and organelle combinations. With the full training data, ConnectoFM achieves a mean Dice of 0.771. When the available labels are reduced to only 20%, the mean Dice remains 0.711, and even at 10% it still reaches 0.673. The degradation is therefore very mild: the absolute drop from full data is only 0.060 at 20% and 0.098 at 10%. In relative terms, ConnectoFM retains 91.0% of its full data performance at 20% labeled data and 84.0% even at 10%. The full low data trajectory is also smooth and stable, with mean Dice values of 0.738, 0.753, and 0.763 at 40%, 60%, and 80% training data, respectively. These results show that once the ConnectoFM representation is available, strong segmentation performance can be obtained from only a small fraction of the original annotation budget.

The contrast with the baselines is substantial. RETINA reaches only 0.369 mean Dice at 20% data and 0.296 at 10%, while PyTC UNet achieves 0.325 at 20% and 0.324 at 10%. Thus, at 10% labeled data, ConnectoFM exceeds RETINA by 0.377 Dice on average and PyTC UNet by 0.349. At 20%, the margins remain similarly large: +0.342 over RETINA and +0.386 over PyTC UNet. These are not isolated improvements on a few favorable settings. At 10% data, ConnectoFM outperforms RETINA on all 22*/*22 matched dataset and organelle tasks and outperforms PyTC UNet on 21*/*22. At 20%, it still wins on 21*/*22 tasks against RETINA and 21*/*22 against PyTC UNet. The same pattern persists throughout the entire data efficiency curve, showing that the advantage of ConnectoFM is systematic rather than anecdotal.

A particularly strong result is obtained when comparing ConnectoFM in the low data regime against baselines trained with the *full* training set. Even with only 10% of the labeled data, ConnectoFM still surpasses RETINA trained on 100% data by an average of 0.241 Dice and performs better on 18*/*22 tasks. At 20% data, the average margin over full data RETINA further increases to 0.279, again with wins on 18*/*22 tasks. ConnectoFM also remains highly competitive against full data PyTC UNet: at 10% labeled data it achieves an average advantage of 0.154 Dice and wins on 12*/*21 tasks, and at 20% labeled data the advantage grows to 0.193 Dice with wins on 14*/*21 tasks. This finding is especially important because it shows that the pretrained connectomics representation does not merely help when all methods are equally label starved. Rather, it enables ConnectoFM to remain competitive even against baselines given *substantially more supervision*.

The superiority of ConnectoFM in low data settings is also visible across multiple organelle categories. At 10% labeled data, ConnectoFM achieves mean Dice scores of 0.692 for membranes, 0.851 for mitochondria, 0.322 for synapses, 0.666 for vesicles, and 0.775 for PSD. The corresponding RETINA scores are 0.358, 0.483, 0.010, 0.079, and 0.009, while PyTC Unet reaches 0.445, 0.409, 0.010, 0.083, and 0.009. The same trend remains at 20% data, where ConnectoFM attains 0.725 for membranes, 0.870 for mitochondria, and 0.371 for synapses, compared with 0.417, 0.670, and 0.010 for RETINA, and 0.439, 0.428, and 0.010 for PyTC UNet. These numbers indicate that the advantage of ConnectoFM is not confined to a single easy organelle class. It extends across both relatively structured categories such as mitochondria and more challenging fine scale targets such as synapses and PSD.

To summarize the entire data efficiency curve in a single number, we also computed the area under the mean Dice versus training fraction curve. ConnectoFM achieves a curve area of 0.743, substantially above RETINA at 0.408 and PyTC UNet at 0.381. This aggregate view confirms the same conclusion reached by the pointwise analysis: ConnectoFM is markedly more data efficient than both baselines. Taken together, these results show that connectomics specific pretraining produces rich and reusable features that remain highly informative even when only 10% to 20% of the labeled training data is available. Because only the segmentation head is trained in this setting, the strong performance directly supports the claim that the frozen ConnectoFM encoder already captures connectomics relevant structure in a form that can be exploited with minimal supervision.

#### 2.6.1 Representation Geometry of ConnectoFM Embeddings Enable Data Efficiency

We already show that ConnectoFM remains effective even when only a small fraction of the labeled training set is available. A natural next question is why this happens. In particular, we want to determine whether the frozen ConnectoFM encoder already organizes the downstream data into a feature space that is easy to separate with a simple decision boundary. If this is true, then strong low-data performance should not require a complex decoder or substantial task-specific adaptation. Instead, even a linear probe trained on top of frozen features should perform well, and the resulting scaling curve should improve smoothly as more labels are added.

To test this directly, we performed a representation geometry and sample complexity analysis on the drosophila-vnc mitochondria dataset. We froze the pretrained ConnectoFM encoder and extracted patch-token embeddings from each image, where each token corresponds to one spatial patch produced by the ViT encoder. Specifically, pixel-level mask annotations were converted to patch-level labels by average-pooling each patch region and assigning a foreground label when the fraction of foreground pixels in that patch exceeded a fixed threshold (0.5). Each token was assigned a binary label, foreground or background, from the corresponding segmentation mask. We then trained only a linear classifier of the form *y* = *Wz*+*b* on top of these frozen embeddings, using training fractions *{*0.1, 0.2, 0.4, 0.6, 0.8, 1.0*}*, and evaluated Dice on the held-out test set.

In addition to linear-probe performance, we explicitly quantify the geometry of the frozen representation. Let *z*_*i*_ ∈ ℝ^*d*^ denote a token embedding with class label *y*_*i*_ ∈ *{*1, …, *C}*. For each class, we compute the class mean 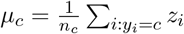, and the global mean 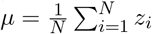. We define the within-class scatter as 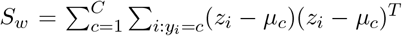 and the between-class scatter as 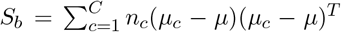. To measure class separability, we compute the optimal Fisher discriminant value 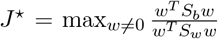, which corresponds to the largest generalized eigenvalue of the matrix pair (*S*_*b*_, *S*_*w*_) [53]. Larger values of *J*^⋆^ are desirable, as they indicate greater linear separability where the distance between class means dominates the variance within the individual classes. We also measure point-wise separation using the margin 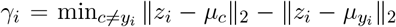. A positive margin means that a token is closer to the center of its true class than to the nearest competing class. Together, these quantities tell us whether the frozen representation already has a favorable geometry for downstream prediction.

The geometry results are summarized in Table 3. Several observations are important. First, the task is highly imbalanced at token level, with only about 5% to 6% foreground tokens across splits. Despite this imbalance, the optimal Fisher discriminant value *J*^⋆^ is consistently greater than 1, indicating that there exist directions in the embedding space where inter-class separation outweighs intra-class spread, thereby confirming the presence of meaningful and exploitable linear structure in the learned representation.. Second, the average margin is consistently positive, with values of 0.5289 on the training split, 0.4710 on validation, and 0.4750 on test. This means that, on average, tokens lie closer to the center of their own class than to the nearest competing class. Third, the fraction of tokens with positive margin is high and remarkably stable, around 88.4% on both validation and test. Taken together, these results show that the frozen ConnectoFM embedding space is not only structured, but also generalizes without an obvious collapse in geometry outside the training split.

**Table 3:**
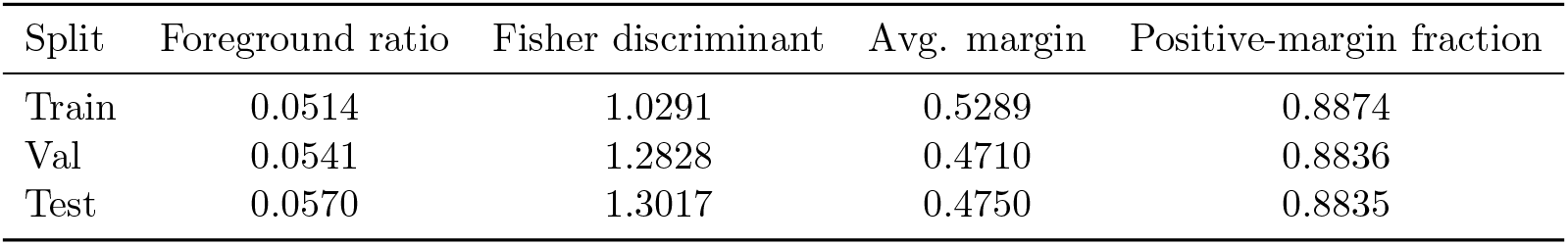
Geometry statistics of the frozen ConnectoFM representation on the drosophila-vnc mitochondria task. Foreground ratio denotes the proportion of foreground tokens. Higher Fisher ratio, larger average margin, and a higher positive-margin fraction indicate better class separation.

| Split | Foreground ratio | Fisher discriminant | Avg. margin | Positive-margin fraction |
| --- | --- | --- | --- | --- |
| Train | 0.0514 | 1.0291 | 0.5289 | 0.8874 |
| Val | 0.0541 | 1.2828 | 0.4710 | 0.8836 |
| Test | 0.0570 | 1.3017 | 0.4750 | 0.8835 |

The linear-probe results in Fig. 6 provide a complementary view of the same phenomenon. Even with only 10% of the labeled tokens, the linear probe reaches a test Dice of 0.5235. As more labels are added, performance improves gradually to 0.5361 at 20%, 0.5358 at 40%, 0.5425 at 60%, 0.5621 at 80%, and 0.5649 at full supervision. Thus, the total gain from 10% to 100% labeled data is only about 0.041 Dice.

**Figure 6:**
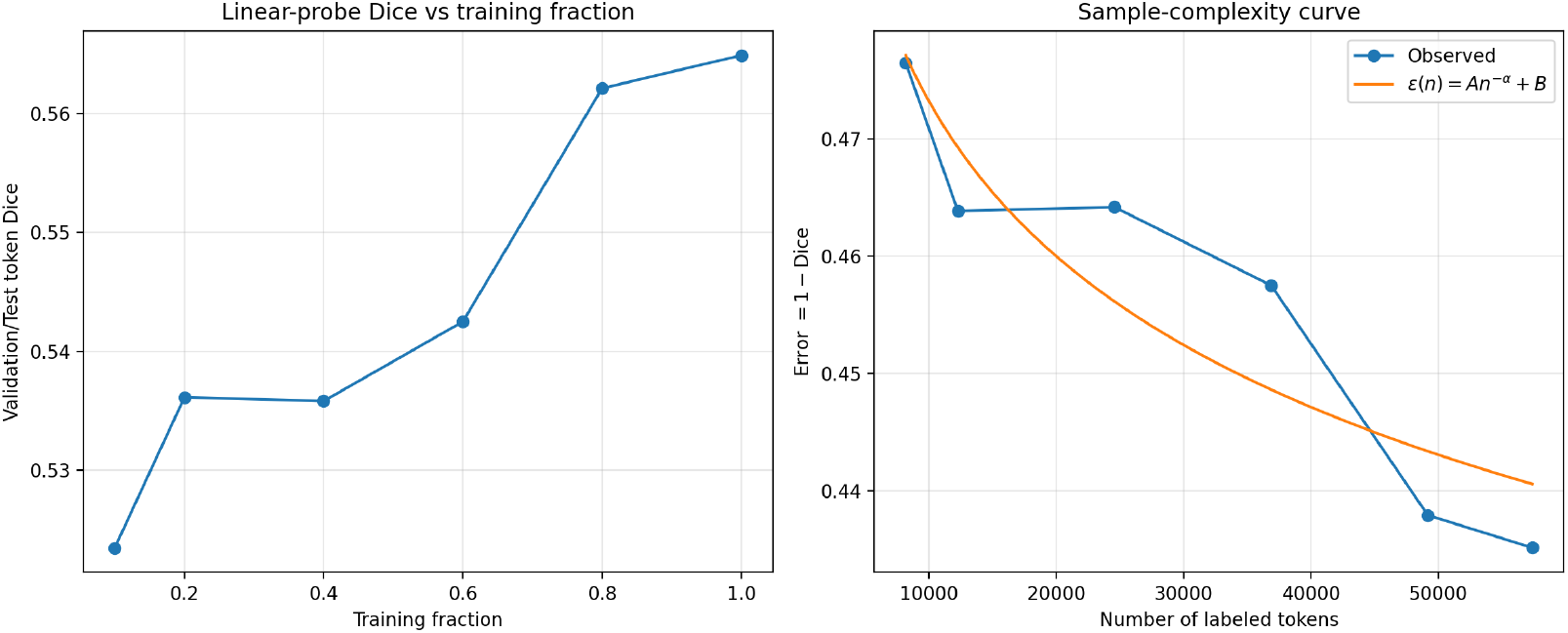
Linear-probe performance and sample complexity of frozen ConnectoFM embeddings on the drosophila-vnc mitochondria dataset. Left: test Dice obtained by training only a linear probe on top of frozen patch-token embeddings at different training fractions. Right: sample-complexity curve for the error *ϵ*(*n*) = 1 − Dice as a function of the number of labeled tokens, together with the fitted power-law model *ϵ*(*n*) = *An*^−*α*^ + *B*.

To quantify this behavior more formally, we fit the error curve *ϵ*(*n*) = *An*^−*α*^ + *B* with *ϵ*(*n*) = 1 − Dice. The fitted parameters are *A* = 0.6905, *α* = 0.0410, and *B* ≈ 0, with RMSE = 0.0062. The small exponent *α* indicates that performance improves steadily but gradually with additional labels, while the very low fitting error shows that the observed scaling behavior is well described by this simple model. The fitted curve therefore supports the same interpretation as the geometry statistics: downstream learning is not starting from a poorly organized embedding space and slowly discovering structure from labels. Rather, the embedding space is already sufficiently aligned with the binary segmentation task, and additional supervision mainly refines an already simple decision boundary.

Overall, this experiment provides a geometric explanation for the data efficiency of ConnectoFM. The Fisher analysis shows that class centers are separated relative to within-class spread, the margin analysis shows that most tokens are already closer to the correct class than to competing classes, and the linear-probe scaling curve shows that only modest additional improvement is obtained as more labels are added. Together, these findings indicate that the frozen ConnectoFM encoder produces a statistically well-structured representation space in which simple linear readouts are already effective, explaining why strong segmentation performance can be achieved with limited supervision.

### 2.7 Beyond 2D: ConnectoFM Retains its Effectiveness in 3D Volumetric Segmentation

We next examined whether the advantage of ConnectoFM extends beyond slice-wise evaluation to volumetric EM segmentation. To examine this, we compared ConnectoFM and RETINA on two 3D mitochondria segmentation benchmarks: Fly Brain Mitochondria Benchmark and Lucchi++. We used a 2D training and inference setup followed by 3D reconstruction. The Lucchi++ dataset provides two annotated volumes of size 165 *×* 1024 *×* 768. We used one full volume for testing and split the other into training and validation subsets in a 90:10 ratio. The Fly Brain Mitochondria benchmark provides a labeled cube of size 256 *×* 256 *×* 256 voxels. We used the first half of the volume for training and validation with a 90:10 split, and the remaining half for testing. In both cases, models were trained and inferred on 2D slices, and the slice-wise predictions were then integrated into volumetric masks using u-Segment3D from the orthogonal slice views [39]. This setting allows us to test whether representations learned from 2D pretraining remain useful when applied within a 3D reconstruction pipeline.

As shown in Table 4, ConnectoFM outperforms RETINA on both 3D benchmarks in overall overlap quality. On Fly Brain Mitochondria Benchmark, ConnectoFM achieves a Dice score of 0.8307 compared with only 0.5706 for RETINA, corresponding to an improvement of 45.58%, and also yields higher IoU, precision, and recall. On Lucchi++, ConnectoFM again performs better overall, with Dice improving from 0.8592 to 0.8833 and IoU from 0.7532 to 0.7910, while maintaining strong precision and comparable recall. The qualitative reconstructions in Fig. 7 further support these trends. In both datasets, ConnectoFM produces volumetric segmentations that more closely follow the annotated mitochondrial morphology, with better continuity and fewer missing structures, whereas RETINA exhibits more fragmented predictions and additional false positive regions. These results show that, although ConnectoFM is pretrained in 2D, its learned representations remain effective in realistic 3D volumetric segmentation workflows.

**Table 4:**
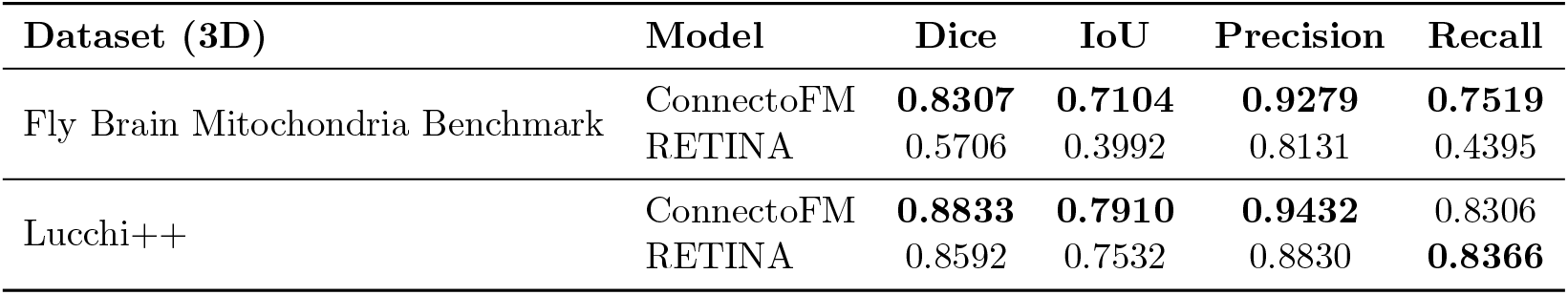
Mitochondria segmentation performance on two 3D EM benchmarks. We report Dice, IoU, precision, and recall for the two foundation models ConnectoFM and RETINA. For each dataset and metric, the better result is shown in bold.

| Dataset (3D) | Model | Dice | IoU | Precision | Recall |
| --- | --- | --- | --- | --- | --- |
| Fly Brain Mitochondria Benchmark | ConnectoFM | **0.8307** | **0.7104** | **0.9279** | **0.7519** |
|  | RETINA | 0.5706 | 0.3992 | 0.8131 | 0.4395 |
| Lucchi++ | ConnectoFM | **0.8833** | **0.7910** | **0.9432** | 0.8306 |
|  | RETINA | 0.8592 | 0.7532 | 0.8830 | **0.8366** |

**Figure 7:**
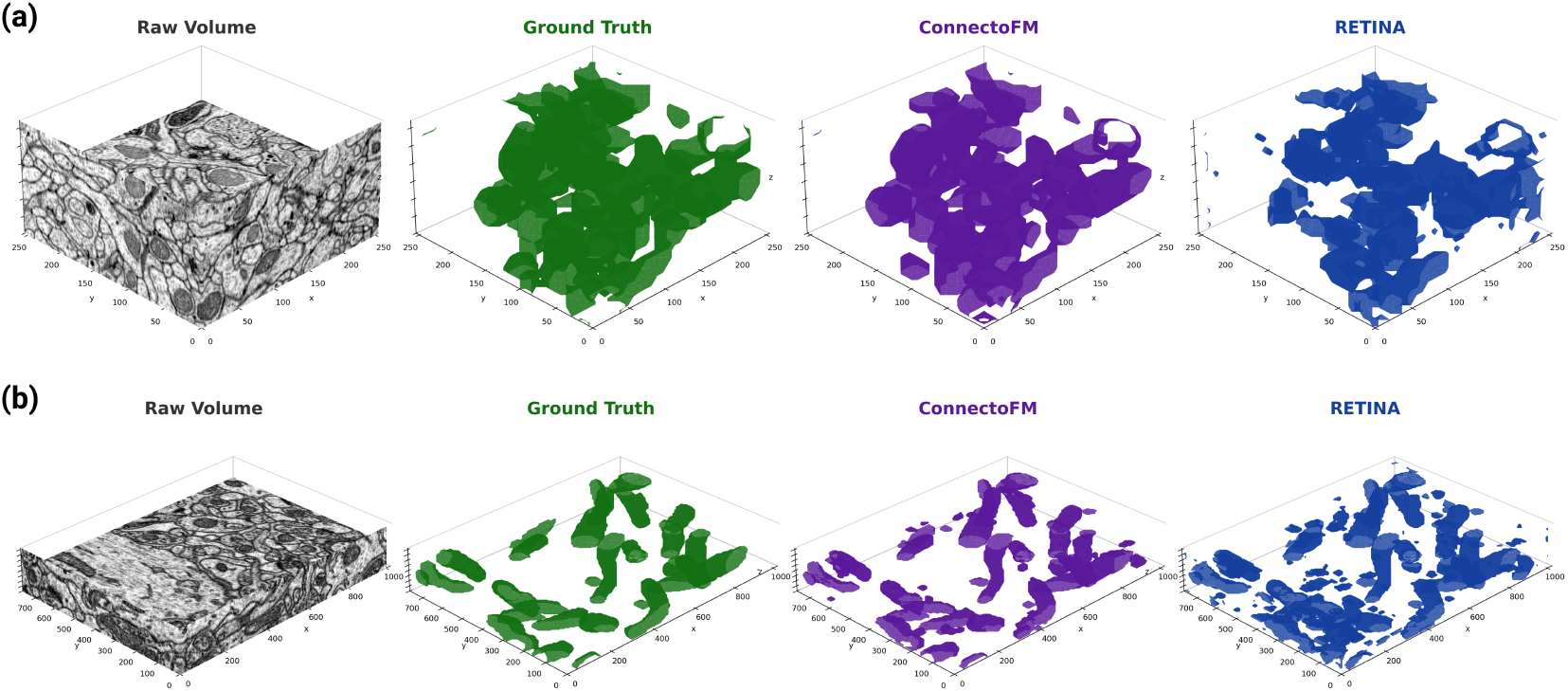
Qualitative comparison of 3D binary segmentation of mitochondria on two EM benchmarks between the two foundation models ConnectoFM and RETINA. **a**, Test volume from Fly Brain Mitochondria Benchmark. **b**, Test volume from Lucchi++. In each case, we show the raw volume, ground truth, ConnectoFM prediction, and RETINA prediction.

## 3 Case Studies

We next examine representative qualitative examples to assess how the predictions of ConnectoFM differ from those of competing methods in practice. Fig. 8 shows one randomly selected example from each of five downstream datasets, spanning binary segmentation, multiclass cell typing, and instance segmentation. Across all three task settings, ConnectoFM produces outputs that more closely match the ground truth, with cleaner boundaries, fewer spurious predictions, and more complete recovery of biologically relevant structures. By contrast, the baseline methods more often yield fragmented, porous, incomplete, or mislabeled outputs. Because microSAM performed substantially worse than the other methods in these examples, we present its qualitative visualizations in the Supplementary Material (see Figure S10).

**Figure 8:**
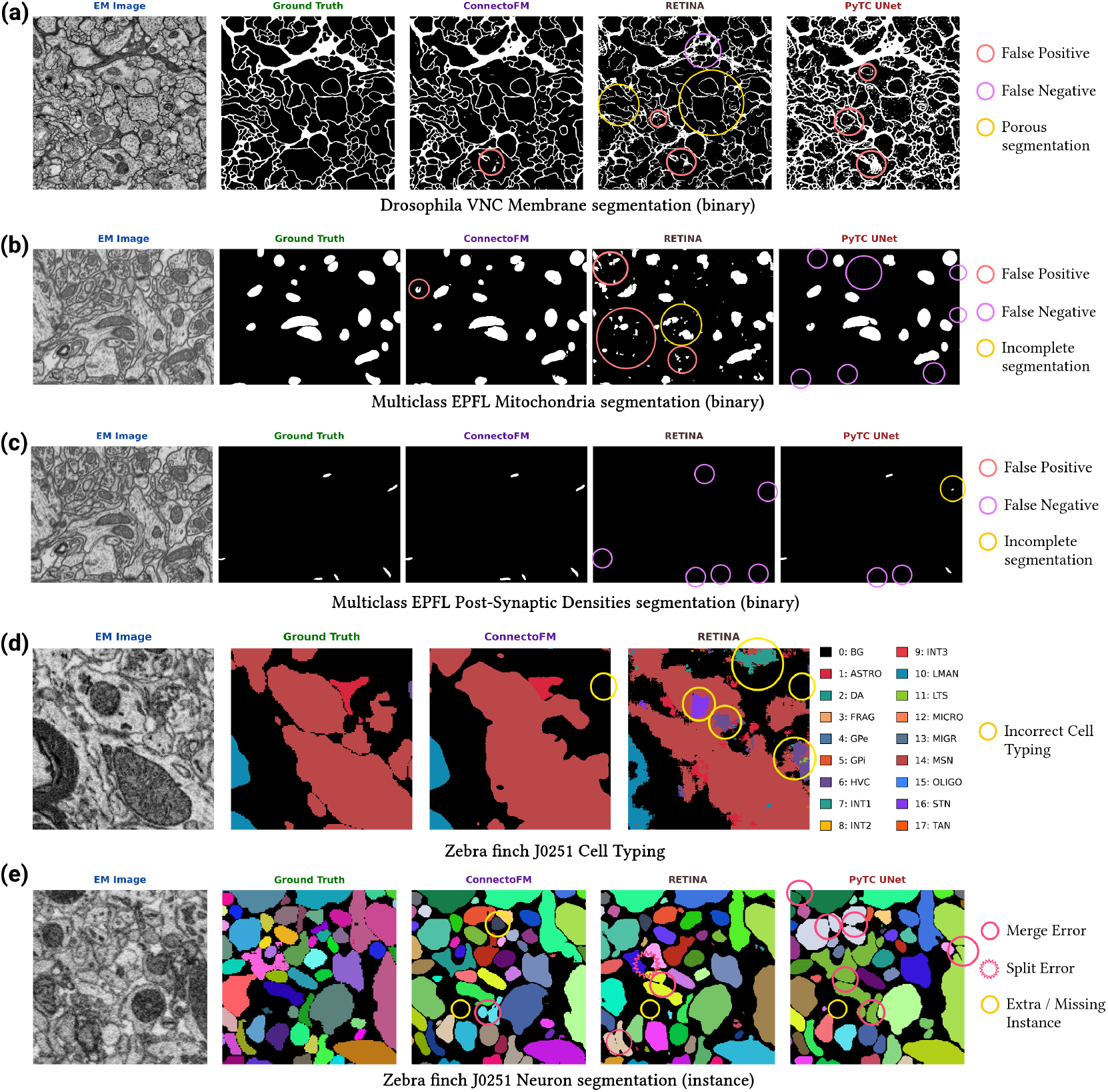
Qualitative comparison of ConnectoFM and baseline methods across downstream connectomics tasks. a–c, Binary segmentation examples comparing ConnectoFM, RETINA, and PyTC UNet, with representative errors highlighted. Across these examples, ConnectoFM produces masks that more closely match the ground truth, whereas the baselines exhibit more false positives, false negatives, and incomplete or porous segmentations. **a**, Membrane segmentation on the *Drosophila* VNC dataset. **b**, Mitochondria segmentation on the Multiclass EPFL dataset. **c**, Post-synaptic dendrite segmentation on the Multiclass EPFL dataset. **d**, Multiclass cell-typing predictions for ConnectoFM and RETINA on the zebra finch J0251 dataset, with representative incorrect assignments highlighted. **e**, Instance segmentation results for ConnectoFM, RETINA, and PyTC UNet on the zebra finch J0251 dataset, with representative merge errors, split errors, and extra or missing instances indicated. microSAM outputs are omitted here for clarity and are provided in the Supplementary Material (see Figure S10). This figure was created with BioRender.com.

In membrane segmentation on the *Drosophila* VNC dataset (Fig. 8a), ConnectoFM produces a largely continuous and non-porous membrane mask that closely follows the ground-truth boundaries, with only a small isolated false-positive region. RETINA, in contrast, exhibits multiple types of error simultaneously, including false positives, false negatives, and visibly porous membrane predictions that disrupt boundary continuity. PyTC UNet recovers much of the large-scale membrane structure, but still introduces several false-positive regions and yields less faithful local delineation than ConnectoFM. This example highlights the ability of ConnectoFM to preserve thin and continuous membrane topology.

The difference remains clear in the Multiclass EPFL examples for mitochondria and post-synaptic densities (PSDs) (Fig. 8b,c). For mitochondria segmentation, ConnectoFM correctly recovers nearly all mitochondrial instances and produces masks that are compact and well aligned with the ground-truth structures, with only one incomplete false-positive instance. RETINA, in contrast, produces fragmented and noisy outputs, with several false-positive regions and incomplete detections, suggesting difficulty in distinguishing true mitochondrial profiles from the surrounding ultrastructure. PyTC UNet detects the major mitochondrial instances, but misses several smaller structures and yields less complete coverage overall.

The PSD example reveals an even sharper contrast. ConnectoFM successfully recovers all PSD instances, preserving their small and spatially sparse morphology despite the difficulty of the task. RETINA fails almost entirely in this example, missing all visible PSD instances even after supervised finetuning. PyTC UNet performs better than RETINA, but still misses multiple PSDs and reconstructs one detected instance only partially. Because PSDs are small, sparse, and directly tied to synaptic organization, these qualitative differences are especially important from a connectomics perspective.

We next consider multiclass cell typing in the Zebra Finch J0251 dataset (Fig. 8d). ConnectoFM produces a cell-type map that is visually much closer to the ground truth, preserving the dominant cellular regions and their boundaries with relatively few mistakes. Its main errors in this example are limited and mostly take the form of small cellular regions predicted as background. RETINA, by contrast, shows several misclassifications across the image. In particular, it confuses portions of the dominant MSN region with STN and introduces additional erroneous labels in areas that should remain background.

Finally, Fig. 8e compares the methods on instance segmentation in the same zebra finch dataset. ConnectoFM produces a faithful instance map overall, with one visible merge error and two localized cases of extra or missing instances. RETINA shows a broader range of failures, including both merge and split errors, together with an additional missing instance. PyTC UNet exhibits several merge errors across neighboring cells and also misses the same instance not recovered by the other methods. Although all methods face challenges in this dense instance-segmentation setting, ConnectoFM remains the most reliable, better preserving object separation and instance completeness.

Taken together, these case studies show that ConnectoFM more consistently preserves continuity, completeness, and semantic correctness across structures that are central to connectome reconstruction.

## 4 Discussion

In this study, we introduced ConnectoFM, the first known foundation model developed specifically for connectomics. By pretraining on 1.7 million unlabeled EM images curated across six species and diverse subdomains, ConnectoFM learns reusable representations that transfer effectively to multiple downstream tasks in connectomics. We show that these representations yield robust performance across binary segmentation, multiclass cell typing, and instance segmentation, while also improving data efficiency in low-label regimes. These results are timely because connectomic reconstruction is becoming increasingly central to neuroscience [54], with broad utility in resolving circuit-level mechanisms of learning and memory [55], linking structure to function [52, 56], enabling large-scale neural modelling [57, 58], and supporting comparative, developmental, and pathological analyses of nervous systems [59–63]. As connectomics datasets continue to expand in scale, diversity, and biological scope, reusable representation learning has become an increasingly important dimension for reliable reconstruction and analysis.

A central finding of our study is that ConnectoFM learns embeddings that reflect biologically meaningful organization in the pretraining corpus rather than merely capturing superficial appearance statistics. In the joint embedding space, samples cluster coherently across species and subdomains, and species-specific projections reveal additional structure aligned with developmental stages, brain regions, cortical layers, and acquisition cohorts. Notably, this organization remains strong even in challenging settings. For example, the eight *C. elegans* developmental subdomains exhibit smoother overlap than other species, consistent with gradual developmental transitions, yet still retain meaningful local neighborhood structure. Conversely, in macaque cortex, ConnectoFM cleanly separates the layer 4 domain from the three layer 2/3 subdomains, indicating sensitivity to fine-grained differences in cortical organization. These patterns are supported quantitatively by strong kNN accuracies in both the joint and species-specific analyses. The reconstruction experiments provide a complementary view of pretraining quality. Despite having access to only 25% of visible patches, ConnectoFM reconstructs augmented EM images with high fidelity across six representative domains, preserving both broad morphology and fine ultrastructural detail. Taken together, these observations suggest that ConnectoFM learns representations that are simultaneously globally coherent and locally informative.

The downstream experiments further show that these pretrained representations are practically useful. Across 22 binary segmentation datasets, ConnectoFM achieves a mean Dice score of 0.771, outperforming a scratch-trained PyTC UNet, RETINA, and microSAM by substantial margins on average. The gains are especially pronounced for structures that are central to connectomics and difficult to segment reliably, including membranes, post-synaptic densities, synapses, and mitochondria. ConnectoFM is also notably effective on small and spatially sparse targets, achieving Dice scores of 0.84 for post-synaptic densities and 0.52 for synapses, where the baseline methods perform poorly overall. These trends are important because small local errors in such structures can have disproportionate effects on downstream topology and inferred connectivity. At the same time, our results also reveal that the benefit of connectomics-specific pretraining is not uniform across all targets: axon segmentation remains a notable exception in which ConnectoFM is competitive with some baselines but does not surpass the best dataset-specific model. This heterogeneity is informative rather than discouraging, as it clarifies where current representations are already strong and where additional structural inductive bias or task-specific adaptation may still be needed.

Beyond binary segmentation, ConnectoFM also shows that a connectomics foundation model can support qualitatively different downstream objectives. In dense multiclass cell typing on the Zebra finch J0251 dataset, ConnectoFM substantially outperforms RETINA across all aggregate metrics while also improving class-wise recognition of both dominant and lower-support cell populations. These gains are particularly valuable because biologically meaningful connectome analysis depends not only on accurate reconstruction of objects, but also on reliable assignment of cellular identity. In instance segmentation, ConnectoFM remains competitive across diverse datasets spanning neurons, axons, mitochondria, and nuclei, despite the additional difficulty of preserving object separation and completeness. Although it does not uniformly outperform all baselines in this setting, its overall competitiveness indicates that the learned representations are general enough to support instance-level reasoning across multiple object scales. More broadly, these results suggest that ConnectoFM captures transferable information that extends beyond a single organelle or supervision type, supporting the central promise of foundation models: one pretrained representation can be reused across multiple tasks, species, and annotation regimes with lightweight downstream adaptation instead of repeated end-to-end retraining from scratch.

This practical utility of ConnectoFM is further reinforced by its strong performance in low-label settings, its extension to 3D volumetric segmentation, and the qualitative evidence provided by the case studies. ConnectoFM remains substantially more data-efficient than the baselines in low-annotation settings, indicating that pretraining on large-scale unlabeled connectomics data reduces the amount of additional task-specific supervision needed for strong downstream performance. This view is further supported by our embedding-geometry analysis, which reveals a well-structured and readily separable feature space, providing an interpretable explanation for ConnectoFM’s strong performance under limited supervision. This is especially important for connectomics, where dense annotation is expensive, proofreading is labor-intensive, and many new datasets are emerging with limited labels. The qualitative examples further clarify how these gains manifest in practice. Compared with RETINA and PyTC UNet, ConnectoFM more consistently produces continuous membrane masks, more complete organelle reconstructions, more faithful PSD recovery, and more accurate cell-type maps, while reducing false positives, false negatives, fragmentation, and label confusions. These visual differences matter because connectomics errors are rarely benign since porous membranes, missed synaptic structures, or incorrect cell-type assignments can propagate into downstream reconstruction and biological interpretation [12, 38, 64–66].

Our study also has several limitations that point to promising directions for future work. First, ConnectoFM is pretrained in 2D rather than natively in 3D. We chose this design because 2D pretraining is considerably more scalable and aligns well with practical connectomics workflows, but fully 3D pretraining could potentially capture volumetric continuity and axial context more directly, at the cost of substantially higher computational budget. Nevertheless, our experiments show that ConnectoFM remains effective in 3D segmentation settings through standard 2D-to-3D reconstruction pipelines, where it consistently outperforms general EM foundation models and produces more coherent volumetric segmentations. Second, while our pretraining corpus is already large and diverse, it still covers a subset of the species, preparations, and imaging protocols represented in the broader connectomics literature. Scaling to larger corpora, more species, and more varied anatomical settings may further improve robustness and transfer. Third, our current framework is built on masked autoencoding with contrastive alignment; other self-supervised objectives, such as DINO- or JEPA-style pretraining [67–70], may offer complementary advantages and deserve systematic evaluation in connectomics. Fourth, although instance segmentation performance is competitive, it does not yet show the same clear margin observed for binary segmentation and cell typing, suggesting that additional architectural refinement or instance-aware adaptation could be beneficial. Relatedly, our downstream adaptation currently relies on frozen features with lightweight task-specific decoders. Future work could explore parameter-efficient finetuning strategies, including low-rank adaptation methods [71, 72], to better study domain adaptation, dataset-specific specialization, and efficient transfer across connectomics settings without requiring full end-to-end retraining. Finally, our study remains limited in interpretability. A deeper analysis of what structures and failure modes are encoded by the representation may help guide both scientific understanding and future model design.

Taken together, our results establish ConnectoFM as a generalizable foundation model for connectomics EM data and show that connectomics-specific pretraining can yield representations that are biologically meaningful, transferable across tasks, and practically useful under realistic annotation constraints. We view this as an important step toward a more scalable computational framework for connectomics, in which large unlabeled EM corpora are leveraged not merely as raw data repositories, but as a basis for reusable representation learning. Future extensions in model scale, data diversity, and pretraining objectives will likely further strengthen this paradigm. More broadly, ConnectoFM suggests that connectomics is ready to move beyond narrowly dataset-specific training toward foundation-model-driven analysis, with the potential to accelerate neural circuit reconstruction and enable richer structure-to-function studies across nervous systems.

## 5 Materials and Methods

### 5.1 Curation of the pretraining corpus

We curated a large and diverse collection of 1.7 million electron microscopy (EM) images spanning 25 distinct domains across 6 species and multiple brain regions. The collection includes the complete nervous system of *C. elegans* [7]; the peripheral nociceptive circuits and ventral nerve cord of *Drosophila melanogaster* [73, 74]; the cerebral cortex of humans (*H. sapiens*) [1]; the primary visual cortex and CA1 hippocampus of mice (*Mus musculus*) [75–78]; the visual cortex of the rhesus macaque (*Macaca mulatta*) [77]; and the whole brain of zebrafish (*Danio rerio*) [79]. Together, these domains capture substantial biological and technical diversity, spanning developmental stages from larval to adult and multiple imaging modalities, including serial section EM (ssEM), serial transmission EM (ssTEM), and serial block-face EM (SBEM). Except for the H01 dataset, which was obtained from its native host, all data were sourced from BossDB. Detailed specifications for each domain are provided in Supplementary Table 1.

### 5.2 Preprocessing of the pretraining corpus

Given that connectomes are stored as massive 3D EM volumes, we first extract random sub-volumes of size 256*×*256*×*256 voxels. However, stochastic sampling often yields cubes dominated by void regions, that is, large contiguous areas of black background pixels devoid of biological structure. To ensure the quality of the pretraining data, we apply a multi-stage thresholding procedure to remove these uninformative volumes. Our filtering scheme uses three parameters: a foreground fraction threshold (*τ*_*f*_ = 0.5), a slice-level threshold (*τ*_*fs*_ = 0.5), and a volume-wide stability threshold (*τ*_*gs*_ = 0.9). We first discard any cube whose total fraction of foreground pixels falls below *τ*_*f*_. We then perform slice-level filtering, where a slice is considered acceptable only if at least *τ*_*fs*_ of its pixels are identified as foreground. A candidate volume is retained only if at least *τ*_*gs*_ of its constituent slices satisfy this criterion. After filtering, the retained 3D volumes are decomposed into 2D images and standardized into 256 *×* 256 patches. This curated collection of unlabeled images constitutes the final pretraining corpus.

### 5.3 Datasets used for downstream evaluation

We evaluated ConnectoFM on a diverse downstream benchmark spanning binary segmentation, multiclass cell typing, and instance segmentation across multiple species, brain regions, and ultrastructural targets. The binary segmentation benchmark comprises 22 dataset–task pairs drawn from 15 datasets, including C. elegans Dauer [63, 80], Drosophila VNC [74], MICrONS Basil [75], FlyWire [3, 81], SNEMI3D [82], CREMI [18], Lucchi++ [15], Kasthuri++ [15], and Multiclass EPFL [83], and covers membranes, mitochondria, mitochondrial boundaries, post-synaptic densities (PSDs), synapses, vesicles, and axons. For multiclass cell typing, we evaluate on the Zebra finch J0251 dataset [84]. For instance segmentation, we evaluate on 7 datasets, including AxonEM-H, AxonEM-M [85], Zebra finch J0126, Zebra finch J0251 [84], CEM-MitoLab 22K [86, 87], NucMM-M, and NucMM-Z [88], covering neuron, axon, mitochondrion, and nucleus instances. Together, these datasets span substantial variation in species, anatomy, and imaging conditions (Supplementary Tables 2 and 3). Although the downstream models operate on 2D image patches, many of these images are drawn from serial-section or volumetric EM datasets and therefore arise naturally within 3D reconstruction workflows. As a result, this evaluation assesses not only slice-level prediction quality, but also the practical usefulness of the learned representations for broader connectomics pipelines in which 2D predictions are integrated across sections for 3D reconstruction.

### 5.4 Pretraining of ConnectoFM

We pretrain ConnectoFM on unlabeled electron microscopy images using masked image modelling with a Vision Transformer [89] based masked autoencoder. Each EM image is resized to 256 *×* 256 prior to augmentation and tokenization. Given an input image *x* ∈ ℝ^*H×W ×*1^ with *H* = *W* = 256, we partition *x* into non overlapping patches of size *P × P* with *P* = 16, producing *N* = (*H/P*)^2^ = 256 patch tokens. Following [90], we randomly mask a fraction (*r* = 0.75) of the patch tokens and retain the remaining (1 − *r*) tokens as visible context. Only the visible patch tokens, together with their positional embeddings, are fed into the encoder to obtain their latent representations. These encoded representations are then concatenated with learnable mask tokens corresponding to the masked patches. The resulting sequence is passed to the decoder, which reconstructs the masked regions. As demonstrated in [90], reconstructing images from such heavily masked inputs encourages the model to learn meaningful and nontrivial representations. This pretraining procedure is illustrated in Fig. 1a.

#### Model architecture

The encoder follows a Vision Transformer Base configuration (ViT-B/16) and is composed of 12 stacked Transformer encoder layers. Each layer contains a multi-head self-attention module and a position-wise feed-forward network, with all layers operating on 768-dimensional token embeddings using 12 attention heads. The decoder adopts a lightweight Vision Transformer Small (ViT-S/16) architecture with 12 stacked Transformer layers. All layers in the decoder operate in a 384-dimensional latent space with 6 attention heads, and are used to reconstruct masked patch tokens from the encoded visible context. After masking, we prepend a learnable `[CLS]` token (with positional embedding) to the visible patch tokens, and process them with the encoder. As shown in Fig. 1a, we then project encoder tokens to the decoder dimension, insert learned mask tokens for missing patches, and apply the decoder with learned positional embeddings. A linear prediction head maps each decoded patch token to a pixel vector of dimension *P* ^2^ · 1.

#### Training objective

We pretrain ConnectoFM with a composite objective that combines masked reconstruction and contrastive alignment to encourage complementary properties in the learned representations. The reconstruction loss forces the encoder to infer missing patch content from sparse context, promoting locality sensitive features that capture fine ultrastructural detail, such as thin membrane boundaries and organelle textures, while also requiring global context to resolve ambiguous regions. However, reconstruction alone can yield representations that are sensitive to nuisance variability introduced by imaging conditions or stochastic preprocessing. To improve invariance and stabilize global semantics, we add a contrastive alignment term that pulls together embeddings of two independently augmented views of the same image and pushes apart embeddings from other images in the batch. This encourages the `[CLS]` representation to encode persistent structural cues while discounting augmentation induced noise, resulting in a representation space that is both structurally descriptive and robust for transfer to downstream connectomics segmentation tasks.

1. **Reconstruction objective**. Let ℳ denote the set of masked patch indices. The decoder predicts a pixel vector 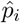 for each patch *i*, and we compute the mean squared error only over masked patches,

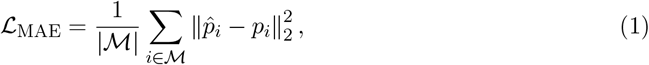

where *p*_*i*_ is the corresponding patch vector obtained by reshaping the input image into patch space.
2. **Contrastive alignment objective**. In addition to masked reconstruction, we impose an embedding alignment term that encourages invariance to stochastic perturbations of the same underlying image. For each input image *x*, we sample two independent augmentations *t*_1_, *t*_2_ ~ 𝒯 and form two views *x*^(1)^ = *t*_1_(*x*) and *x*^(2)^ = *t*_2_(*x*). Let *f*_*θ*_ denote the encoder and let *h*^(*v*)^ = *f*_*θ*_(*x*^(*v*)^) ∈ ℝ^*d*^ be the representation extracted from the encoder `[CLS]` token. We apply *ℓ*_2_ normalization, *z*^(*v*)^ = *h*^(*v*)^*/*∥*h*^(*v*)^∥_2_, and define the temperature scaled similarity, 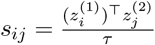. Using the effective batch of size *B* obtained by aggregating representations across all distributed workers, we optimize a symmetric InfoNCE objective

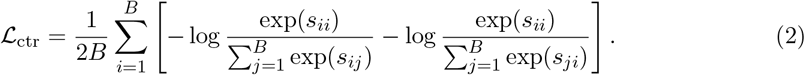

The overall pretraining objective combines reconstruction and contrastive alignment,

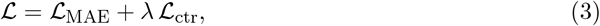

where *λ* controls the strength of the alignment term. The augmentation distribution *𝒯* follows an MAE style design. We apply random resized cropping to the target resolution, sampling the crop area from the scale range [0.2, 1.0] and using bicubic interpolation, followed by random horizontal flipping with probability 0.5. The resulting view is converted to a tensor and standardized using fixed channel wise mean and standard deviation. When forming the contrastive term, two independent draws from *𝒯* are applied to the same image to generate paired views.

### 5.5 Downstream adaptation

We evaluate ConnectoFM by transferring its pretrained encoder to supervised downstream prediction tasks. The ConnectoFM encoder remains frozen and only a lightweight task-specific decoder is trained. Given an input image *x* ∈ ℝ^*H×W*^, we first partition *x* into non-overlapping crops of size 256 *×* 256 and apply the same normalization used during pretraining. The normalized input is then processed by the ConnectoFM encoder *f*_*θ*_, which produces a sequence of token embeddings

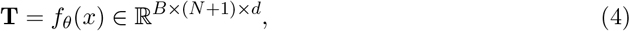

where *d* = 768 and *N* = (*H/P*)^2^ for patch size *P* = 16. We discard the `[CLS]` token and retain the patch tokens **T**_patch_ ∈ ℝ^*B×N×d*^ as dense representations for downstream decoding. For all tasks, training is performed on fixed-size crops extracted on a regular grid. During training, we apply label-preserving geometric augmentations, including random horizontal and vertical flips and random rotations by multiples of 90^°^. At inference time, full-resolution images are processed using an overlapping sliding-window strategy, and predictions from overlapping windows are averaged to reduce boundary artifacts.

#### Binary segmentation

For binary segmentation, we attach a lightweight convolutional decoder to the frozen encoder. Patch tokens are first projected to a lower-dimensional channel space, reshaped into a coarse spatial grid, and then progressively upsampled through a sequence of learned transposed-convolution blocks with interleaved normalization and nonlinearities until full image resolution is recovered. A final 1 *×* 1 convolution followed by a sigmoid activation produces a dense foreground probability map *ŷ* ∈ [0, 1]^*H×W*^. The decoder is trained using a composite loss consisting of pixel-wise binary cross-entropy and Dice loss. Optimization is performed with AdamW and a cosine annealing learning-rate schedule.

#### Cell typing

For cell typing, we cast the task as multiclass pixel classification. The frozen ConnectoFM encoder is paired with a lightweight multiclass decoder that outputs per-pixel logits over cell types. The model is trained with a weighted cross-entropy loss to account for class imbalance, where class weights are computed from the empirical pixel frequencies in the training set. Final predictions are obtained by averaging logits over overlapping sliding-window crops and assigning each pixel to the class with maximal predicted probability.

#### Instance segmentation

For instance segmentation, we adopt an affinity-based formulation. Using the same frozen-encoder adaptation as in binary segmentation, the task-specific decoder predicts a multi-channel affinity map, where each channel encodes pixel connectivity for a predefined spatial offset. Ground-truth affinities are derived directly from the instance labels, and the model is trained with the same binary cross-entropy plus Dice objective applied channel-wise. During inference, overlapping affinity predictions are averaged and converted to instance masks by thresholding followed by connected-component extraction.

### 5.6 Implementation details

ConnectoFM is trained using the AdamW optimizer with a learning rate of 1.5*×*10^−4^ and weight decay 0.05. Pretraining is carried out for 30 epochs with a per device batch size of 48, resulting in an effective global batch size of 192 across 4 Nvidia RTX 4090 GPUs. In total, pretraining required approximately 48 hours. All experiments are initialized with a fixed random seed (42) to ensure reproducibility. Training is performed using PyTorch *DistributedDataParallel* with the NCCL backend. To enable stable and efficient large-scale training, we employ automatic mixed precision with dynamic gradient scaling, and clip gradients to a maximum global norm of 3.0. To guarantee deterministic data ordering across distributed workers and support fault tolerant training, we precompute global batch schedules for all epochs and shard them consistently across devices. This design allows training to be resumed from an arbitrary global batch within an epoch without altering the effective data distribution. Model checkpoints include the encoder and decoder parameters, optimizer state, mixed precision scaler state when applicable, and all relevant random number generator states. 4% of the pretraining corpus (68,401 images) is held out to evaluate the domain organization and reconstruction capability of ConnectoFM representations. For downstream evaluation, we freeze the ConnectoFM encoder (86M parameters) and train a lightweight convolutional decoder head (1.88M parameters) on top of the frozen features. Each downstream dataset is partitioned into training, validation, and test splits with a ratio of 70:15:15. Training is performed for 200 epochs with a batch size of 16. All training for downstream adaptation are performed in one Nvidia RTX 4090 GPU with a fixed seed of 42. Separate models are trained for binary segmentation, instance segmentation, and cell typing. To reduce overfitting, we apply early stopping with a patience of 50 epochs.

## Supporting information

Supplementary Material with additional tables and figures

