## Supplementary Material with additional tables and figures for "ConnectoFM: A Foundation Model for Learning the Language of the Connectome"

This supplementary material presents details on the datasets, evaluation metrics, and additional tables and figures.

#### 1 Evaluation Metrics

We briefly describe the evaluation metrics used across the three downstream tasks.

##### 1.1 Binary Segmentation

For binary segmentation, we report four standard pixel-level metrics computed between the predicted mask $P$ and the ground-truth mask $G$.

**Dice Score.** The Dice score (equivalently, the F1 score for binary segmentation) measures the overlap between $P$ and $G$:

$$\mathrm{Dice}(P, G) = \frac{2|P \cap G|}{|P| + |G|} = \frac{2\,\mathrm{TP}}{2\,\mathrm{TP} + \mathrm{FP} + \mathrm{FN}}, \tag{S1}$$

where TP, FP, and FN denote true positives, false positives, and false negatives, respectively. A score of 1 indicates perfect overlap.

**Intersection over Union (IoU).** Also known as the Jaccard index, IoU measures the fraction of the union covered by the intersection:

$$\mathrm{IoU}(P, G) = \frac{|P \cap G|}{|P \cup G|} = \frac{\mathrm{TP}}{\mathrm{TP} + \mathrm{FP} + \mathrm{FN}}. \tag{S2}$$

**Precision and Recall.** Precision quantifies the proportion of predicted positives that are correct, while recall measures the fraction of true positives that are recovered:

$$\mathrm{Precision} = \frac{\mathrm{TP}}{\mathrm{TP} + \mathrm{FP}}, \qquad \mathrm{Recall} = \frac{\mathrm{TP}}{\mathrm{TP} + \mathrm{FN}}. \tag{S3}$$

High precision indicates few false positives (minimal over-segmentation), while high recall indicates few false negatives (minimal under-segmentation).

##### 1.2 Cell Typing

For multiclass cell-type classification, we report pixel-level accuracy together with macro F1, weighted F1, and mean IoU. Let $C$ denote the number of cell-type classes and let $\mathrm{TP}_c$, $\mathrm{FP}_c$, $\mathrm{FN}_c$ be the per-class counts.

**Pixel Accuracy.** The fraction of pixels assigned the correct class label:

$$\text{Accuracy} = \frac{\sum_{c=1}^{C} \text{TP}_c}{\text{total pixels}}. \tag{S4}$$

**Macro F1.** The unweighted mean of per-class F1 scores, giving equal importance to each class regardless of its frequency:

$$\text{Macro F1} = \frac{1}{C} \sum_{c=1}^{C} \frac{2\,\text{TP}_c}{2\,\text{TP}_c + \text{FP}_c + \text{FN}_c}. \tag{S5}$$

**Weighted F1.** The support-weighted mean of per-class F1 scores, which accounts for class imbalance:

$$\text{Weighted F1} = \sum_{c=1}^{C} w_c \cdot \frac{2\,\text{TP}_c}{2\,\text{TP}_c + \text{FP}_c + \text{FN}_c}, \tag{S6}$$

where $w_c$ is the proportion of ground-truth pixels belonging to class $c$.

**Mean IoU (mIoU).** The unweighted mean of per-class IoU scores:

$$\text{mIoU} = \frac{1}{C} \sum_{c=1}^{C} \frac{\text{TP}_c}{\text{TP}_c + \text{FP}_c + \text{FN}_c}. \tag{S7}$$

##### 1.3 Instance Segmentation

For instance segmentation, we report metrics based on information theory and cluster agreement, following standard practice in connectomics evaluation.

**Variation of Information (VOI).** VOI quantifies the distance between two segmentations $P$ and $G$ as the sum of their conditional entropies:

$$\text{VOI}(P, G) = H(P \mid G) + H(G \mid P), \tag{S8}$$

where $H(P \mid G)$ is the VOI-split term (measuring over-segmentation, i.e., ground-truth segments split across multiple predicted segments) and $H(G \mid P)$ is the VOI-merge term (measuring under-segmentation, i.e., multiple ground-truth segments merged into one predicted segment). We report both terms individually as well as their sum, VOI-total. Lower values indicate better segmentation.

**Normalized Variation of Information (NVI).** NVI normalizes VOI by the joint entropy $H(P, G)$ to produce a scale-invariant measure:

$$\text{NVI}(P, G) = \frac{\text{VOI}(P, G)}{H(P, G)} = 1 - \frac{I(P, G)}{H(P, G)}, \tag{S9}$$

where $I(P, G)$ is the mutual information between $P$ and $G$. NVI ranges in $[0, 1]$ and is less sensitive to the absolute number of segments, making it particularly suitable for comparing across datasets of varying scale. Lower NVI indicates better performance.

**Rand F-score.** The Rand F-score is derived from the Rand index and measures pairwise pixel agreement between $P$ and $G$. For each pair of pixels, a prediction is correct if both pixels are assigned to the same segment in $P$ and $G$, or to different segments in both. The Rand F-score is the harmonic mean of Rand precision and Rand recall computed over all pixel pairs. Higher values indicate better instance-level agreement.

### 2 Supplementary Tables

**Table S1:** Species, region, life stage, and microscopy types of the data domains used in the pretraining corpus of ConnectoFM

| **Domain** | **Species** | **Region** | **Life Stage** | **Microscopy Type** |
|---|---|---|---|---|
| C. elegans – L1 newborn | *C. elegans* | Whole nervous system | First larval (immediately after birth) | ssEM |
| C. elegans – L1 early hours | *C. elegans* | Whole nervous system | First larval (few hours after birth) | ssEM |
| C. elegans – Mid L2 stage | *C. elegans* | Whole nervous system | Second larval (midway) | ssEM |
| C. elegans – Late L1 stage | *C. elegans* | Whole nervous system | Shortly before second larval | ssEM |
| C. elegans – L2 stage | *C. elegans* | Whole nervous system | Second larval | ssEM |
| C. elegans – L3 stage | *C. elegans* | Whole nervous system | Third larval | ssEM |
| C. elegans – Young adult | *C. elegans* | Whole nervous system | Young adult | ssEM |
| C. elegans – Adult | *C. elegans* | Whole nervous system | Young adult | ssEM |
| Drosophila – L3 larva | Drosophila | Peripheral nociceptive circuit (larval) | Third instar larva | TEM |
| Drosophila – Ventral nerve cord | Drosophila | Ventral nerve cord | Adult (female) | TEM |
| Human – Cortex (H01) | Human | Temporal lobe cerebral cortex | Adult | SEM |
| Mouse – Visual Cortex (MICrONS Basil) | Mouse | Primary visual cortex (6 cortical layers) | Postnatal day 87 | Serial EM |
| Mouse – Visual Cortex (MICrONS Minnie) | Mouse | Primary visual cortex (6 cortical layers) | Postnatal day 87 | Serial EM |
| Mouse – CA1 hippocampus | Mouse | Distal dendrites of CA1 hippocampus pyramidal region | Adult (10–14 weeks) | ssTEM |

| Domain | Species | Region | Life Stage | Microscopy Type |
|---|---|---|---|---|
| Mouse – Layer 2/3 Day 105 | Mouse | Primary visual cortex, layer 2/3 | Postnatal day 105 | Serial EM |
| Mouse – Layer 2/3 Day 523 | Mouse | Primary visual cortex, layer 2/3 | Postnatal day 523 | Serial EM |
| Mouse – Layer 4 Day 6 | Mouse | Primary visual cortex, layer 4 | Postnatal day 6 | Serial EM |
| Mouse – Layer 4 Day 105 | Mouse | Primary visual cortex, layer 4 | Postnatal day 105 | Serial EM |
| Mouse – Layer 4 Day 523 | Mouse | Primary visual cortex, layer 4 | Postnatal day 523 | Serial EM |
| Mouse – Cerebellum (Day 56) | Mouse | Cerebellar molecular layer | Postnatal day 56 | SBEM |
| Macaque – Layer 2/3 Day 3000 | Macaque | Primary visual cortex, cortical layers | Postnatal day 3000 | Serial EM |
| Macaque – Layer 2/3 Day 7 | Macaque | Primary visual cortex, cortical layers | Postnatal day 7 | Serial EM |
| Macaque – Layer 2/3 Day 75 | Macaque | Primary visual cortex, cortical layers | Postnatal day 75 | Serial EM |
| Macaque – Layer 4 Day 3000 | Macaque | Primary visual cortex, cortical layers | Postnatal day 3000 | Serial EM |
| Zebrafish – Whole brain | Zebrafish | Whole brain | 5.5 days post fertilization | ssEM |

**Table S2:** Summary of the binary segmentation datasets used in this study

| **Domain** | **Species** | **Brain Region** | **Organelle(s)** | **Image Count** | **Image Size** |
|---|---|---|---|---|---|
| C. elegans Dauer | *C. elegans* | Whole nervous system | Membrane | 48 | $2983 \times 3001$ |
| Mouse Cerebellum (P14) | Mouse | Cerebellar cortex | Membrane | 14 | $4097 \times 4097$ |
| FlyWire | Drosophila | Whole brain | Membrane | 1025 | $1024 \times 1024$ |
| Berghia Connective | Berghia | Rhinophore ganglion region | Membrane | 18 | $2253 \times 1941$ |
| MICrONS Basil | Mouse | Visual cortex | Membrane | 1232 | $1024 \times 1024$ |
| Octopus Glia | Octopus | Vertical lobe deep neuropil | Membrane | 22 | $4529 \times 2379$ |
| Octopus SFL tract | Octopus | Superior frontal lobe tract / vertical lobe | Membrane | 40 | $2265 \times 1189$ |
| SNEMI3D | Mouse | Cortex | Membrane | 100 | $1024 \times 1024$ |
| Mouse Whole brain | Mouse | Whole brain | Membrane | 339 | $771 \times 771$ |
| Fly brain benchmark | Drosophila | Adult fly brain | Mitochondria | 256 | $255 \times 255$ |
| Kasthuri++ | Mouse | Primary somatosensory cortex | Mitochondria | 160 | $1463 \times 1613$ |
| Lucchi++ | Mouse | CA1 hippocampus | Mitochondria | 330 | $1024 \times 768$ |
| Drosophila VNC | Drosophila | Ventral nerve cord | Membrane, mitochondria, synapses | 20 | $1024 \times 1024$ |
| Multiclass EPFL | Mouse | CA1 hippocampus | Membrane, mitochondria, PSD, axon, vesicles | 68 | $1024 \times 768$ |
| Drosophila – CREMI | Drosophila | Adult brain | Synapses | 375 | $1250 \times 1250$ |

**Table S3:** Summary of the instance segmentation datasets used in this study

| **Domain** | **Species** | **Brain Region** | **Organelle(s)** | **Image Count** | **Image Size** |
|---|---|---|---|---|---|
| AxonEM-H | Human | Cerebral cortex | Axon | 810 | $1536 \times 1536$ |
| AxonEM-M | Mouse | Cerebral cortex | Axon | 810 | $1536 \times 1536$ |
| Zebra finch J0126 | Zebra finch | Area X (basal ganglia song system) | Neuron | 4862 | $256 \times 256$ |
| Zebra finch J0251 | Zebra finch | Area X (song system) | Neuron | 5120 | $256 \times 256$ |
| CEM-MitoLab 22K | Mixed | Mixed | Mitochondrion | 21871 | $224 \times 191$ |
| NucMM-M | Mouse | Visual cortex | Nucleus | 1536 | $192 \times 192$ |
| NucMM-Z | Zebrafish | Nearly whole brain | Nucleus | 3456 | $64 \times 64$ |

**Table S4: Average inference time per sample across downstream tasks.** We report the average inference time per sample in seconds for each task.

| **Downstream Task** | **Avg. inference time / sample (s)** |
|---|---|
| Binary segmentation | 0.571 |
| Cell typing | 0.047 |
| Instance segmentation | 0.374 |

### 3 Supplementary Figures

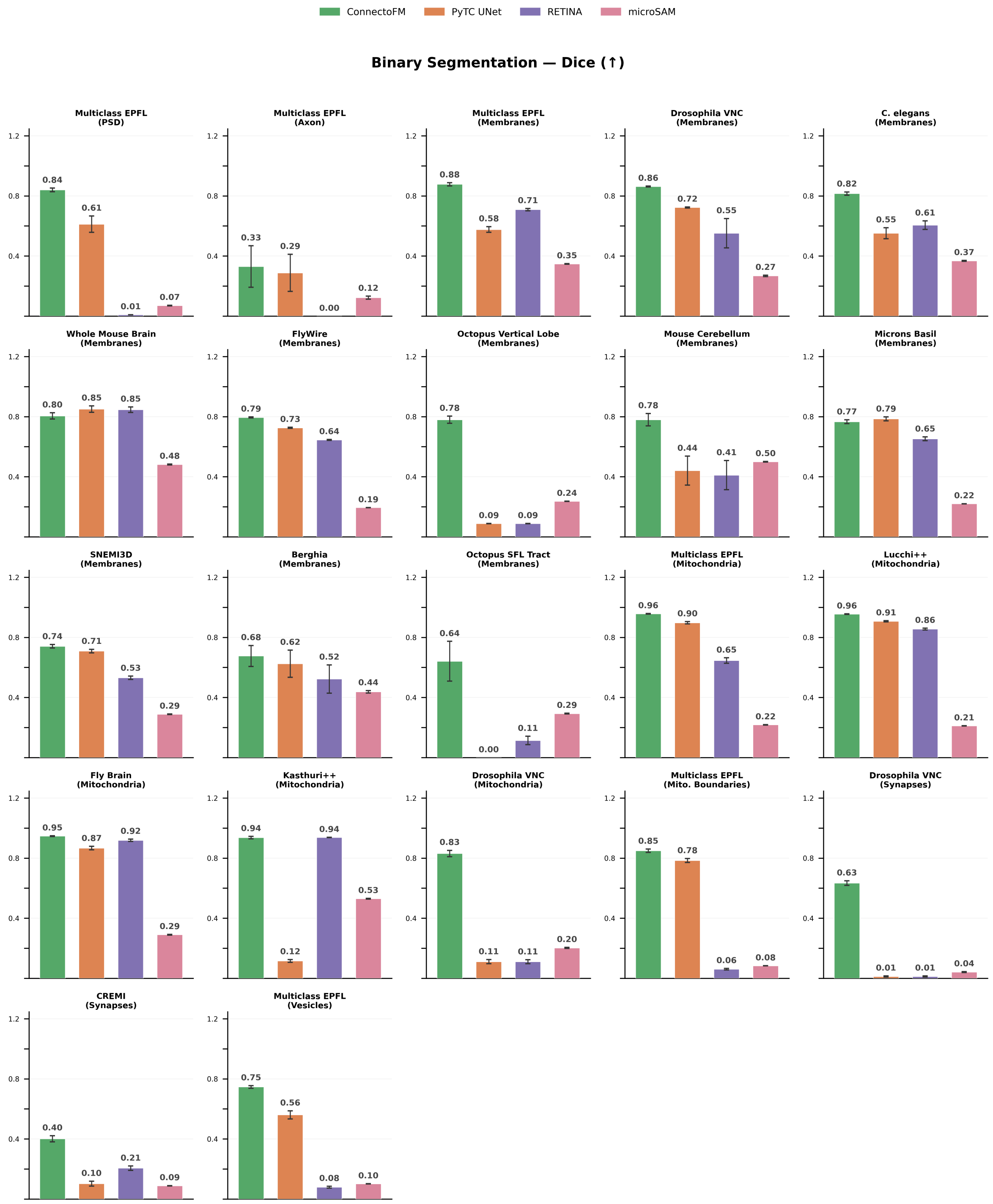


**Figure S1:** Comparison of Dice scores among ConnectoFM, PyTC UNet, RETINA, and microSAM in binary segmentation.

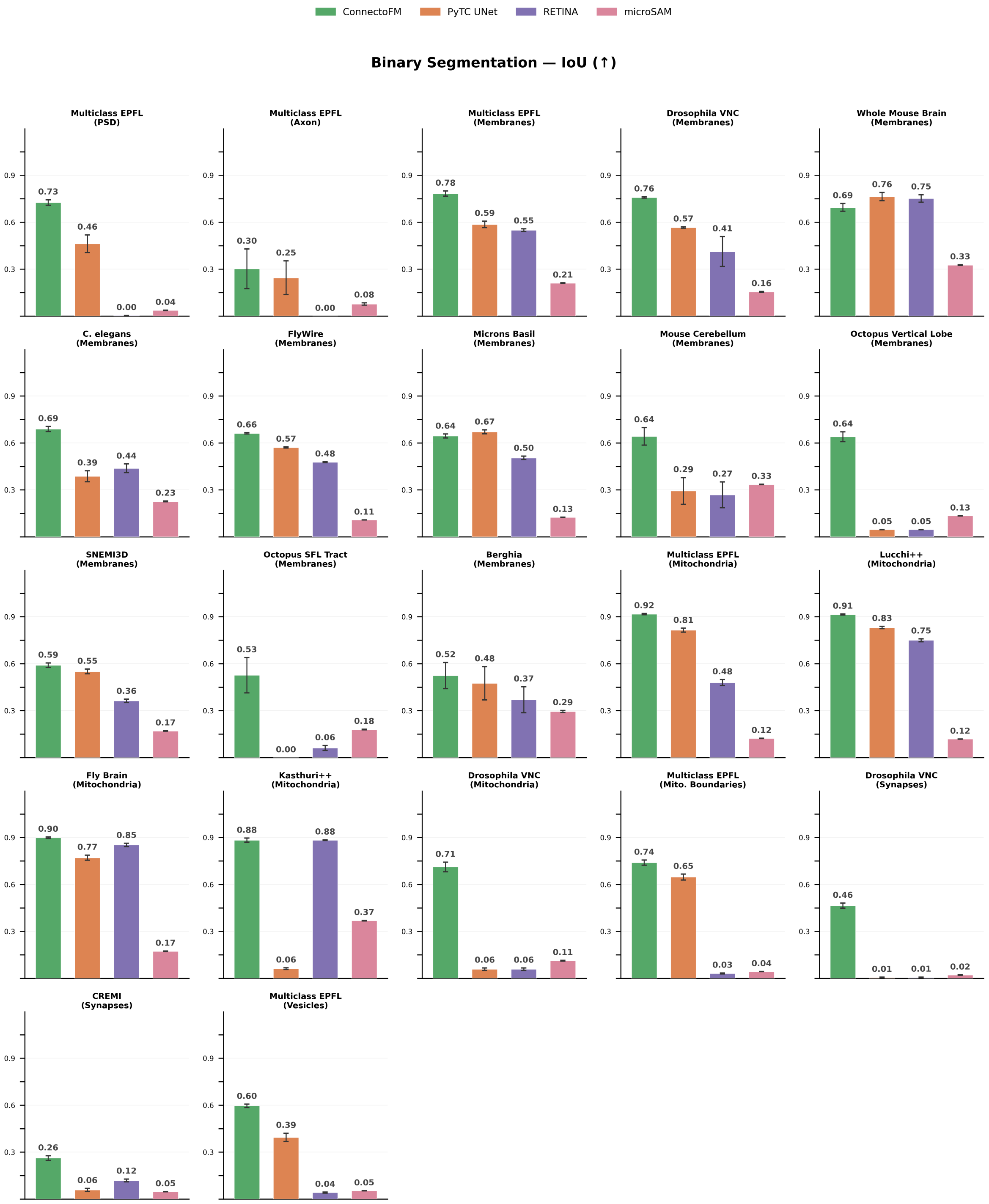


**Figure S2:** Comparison of IoU scores among ConnectoFM, PyTC UNet, RETINA, and microSAM in binary segmentation.

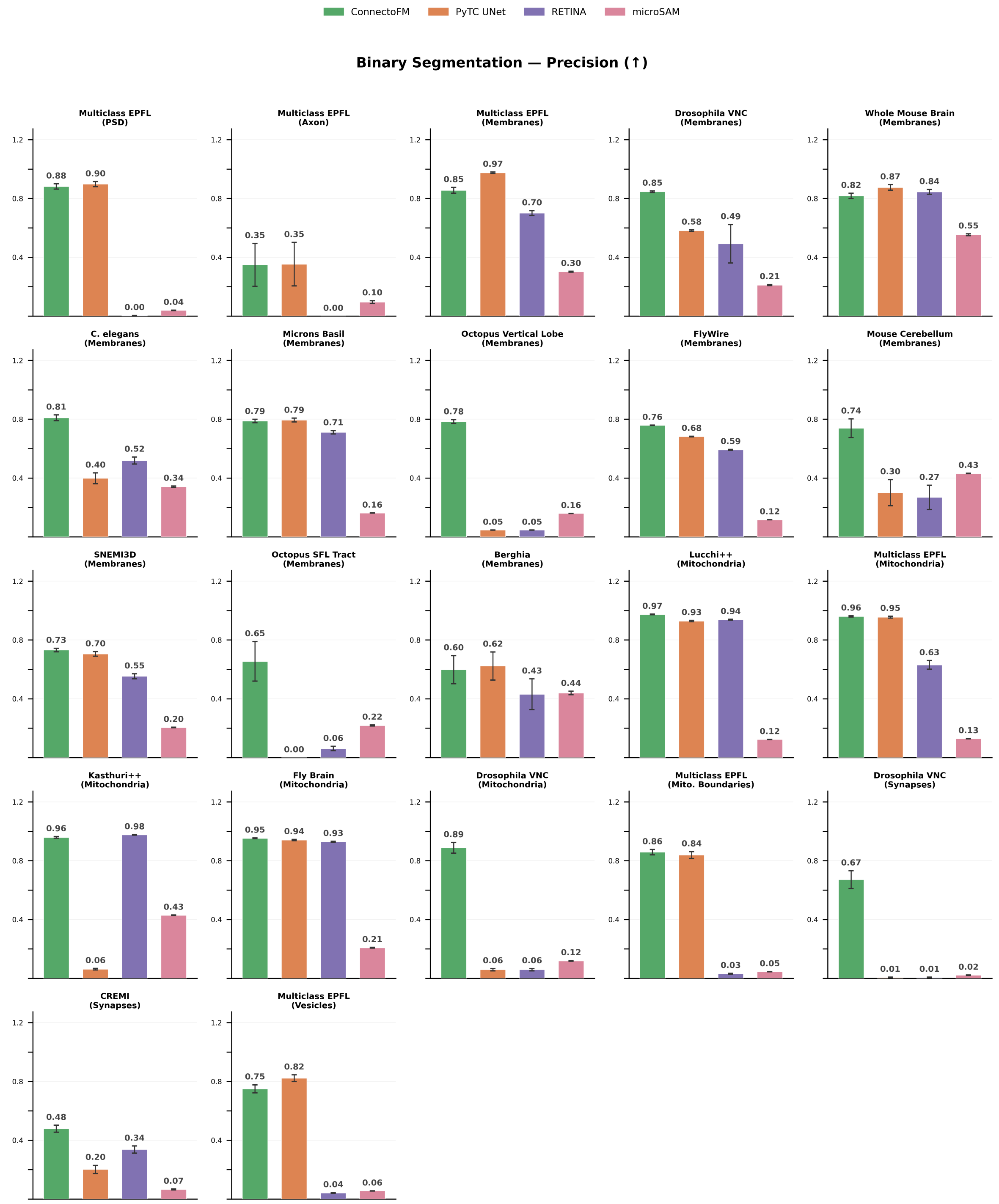


**Figure S3:** Comparison of Precision scores among ConnectoFM, PyTC UNet, RETINA, and microSAM in binary segmentation.

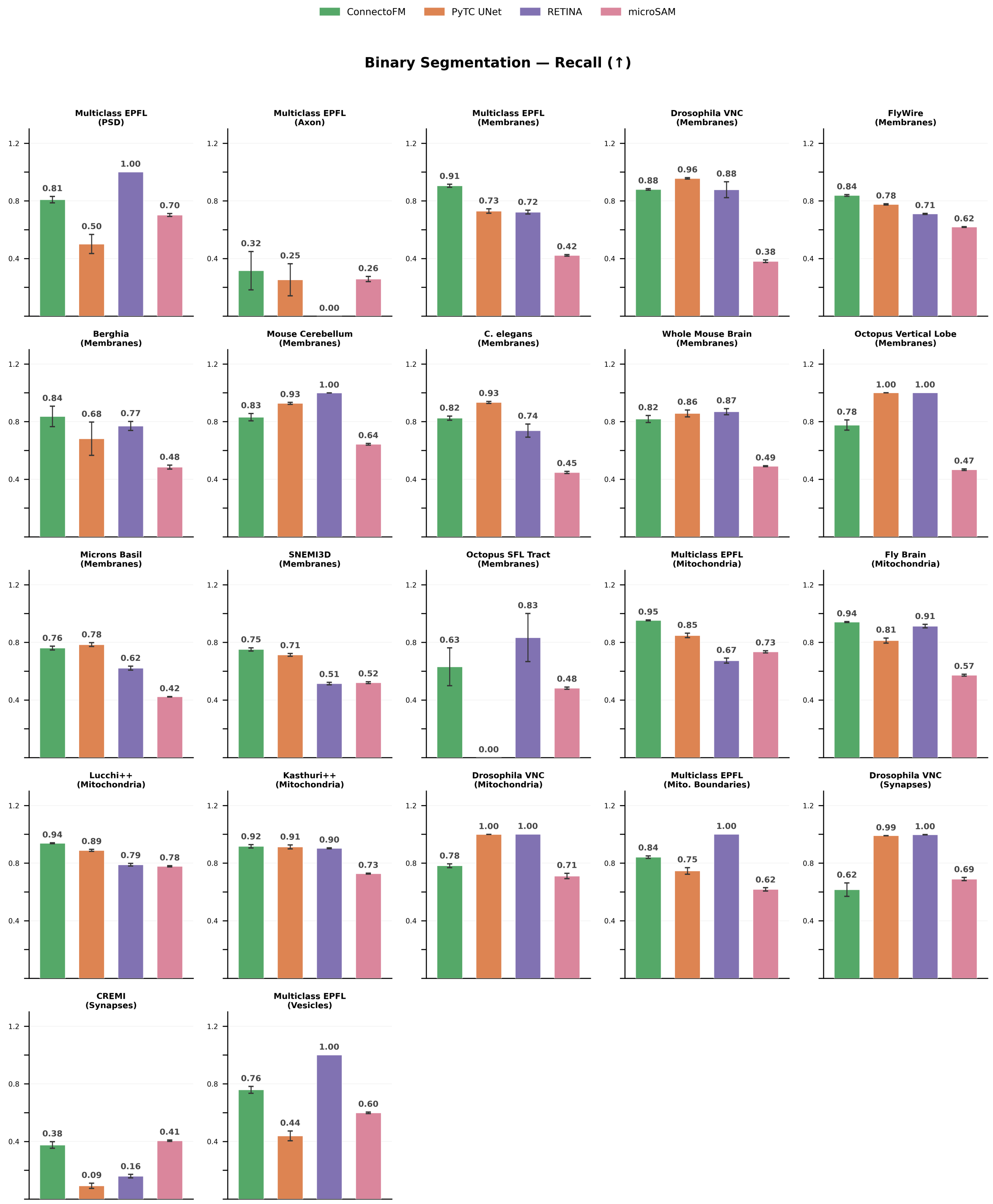


**Figure S4:** Comparison of Recall scores among ConnectoFM, PyTC UNet, RETINA, and microSAM in binary segmentation.

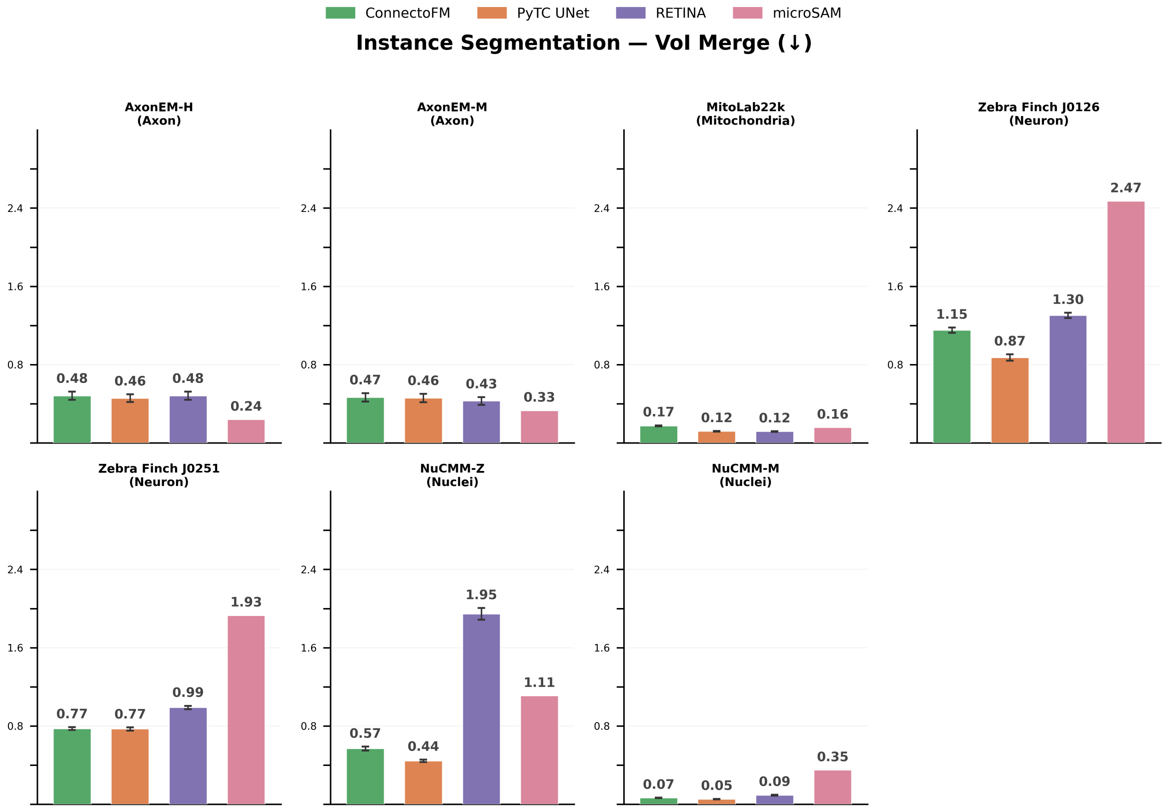


**Figure S5:** Comparison of VOI Merge scores among ConnectoFM, PyTC UNet, RETINA, and microSAM in instance segmentation.

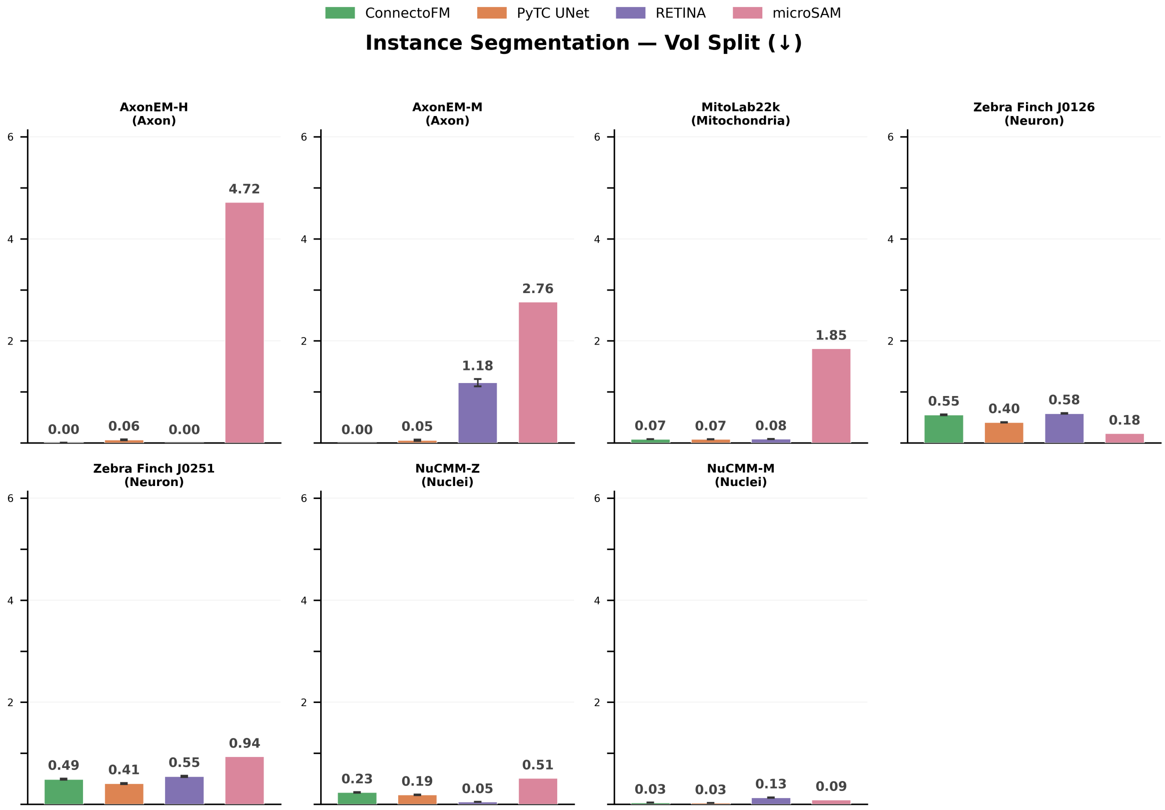


**Figure S6:** Comparison of VOI Split scores among ConnectoFM, PyTC UNet, RETINA, and microSAM in instance segmentation.

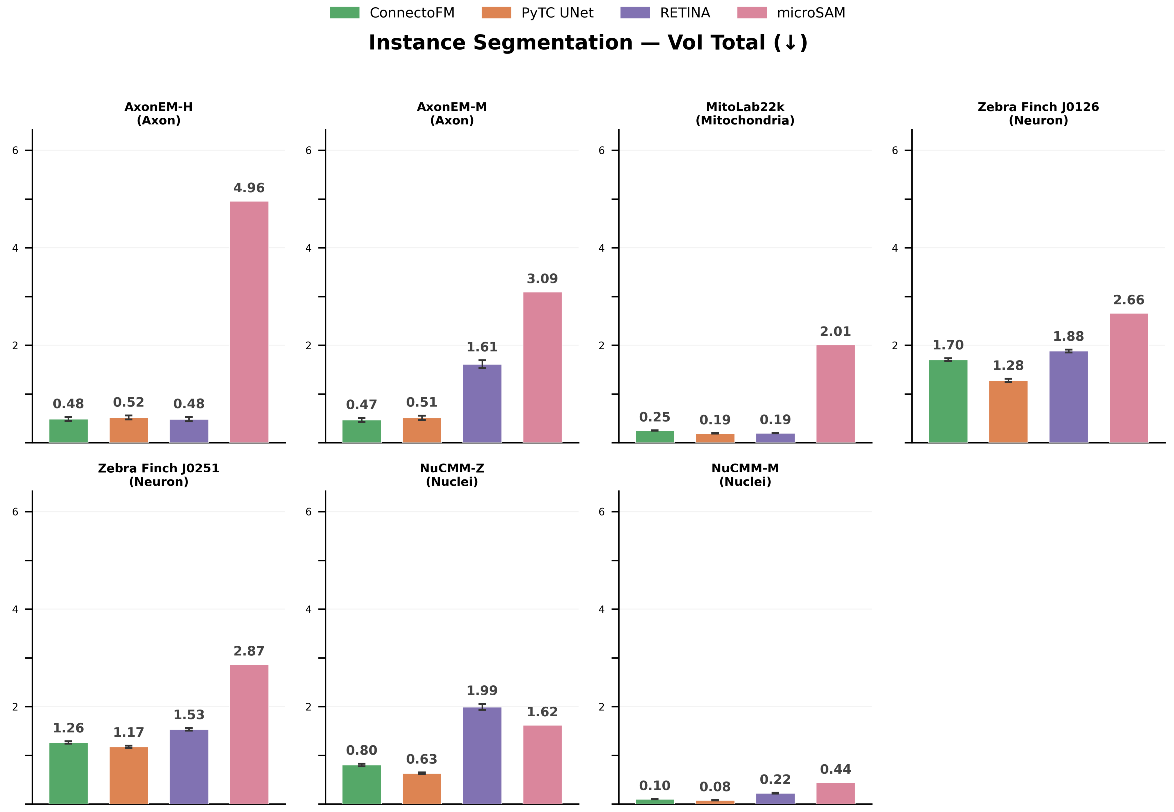


**Figure S7:** Comparison of VOI Total scores among ConnectoFM, PyTC UNet, RETINA, and microSAM in instance segmentation.

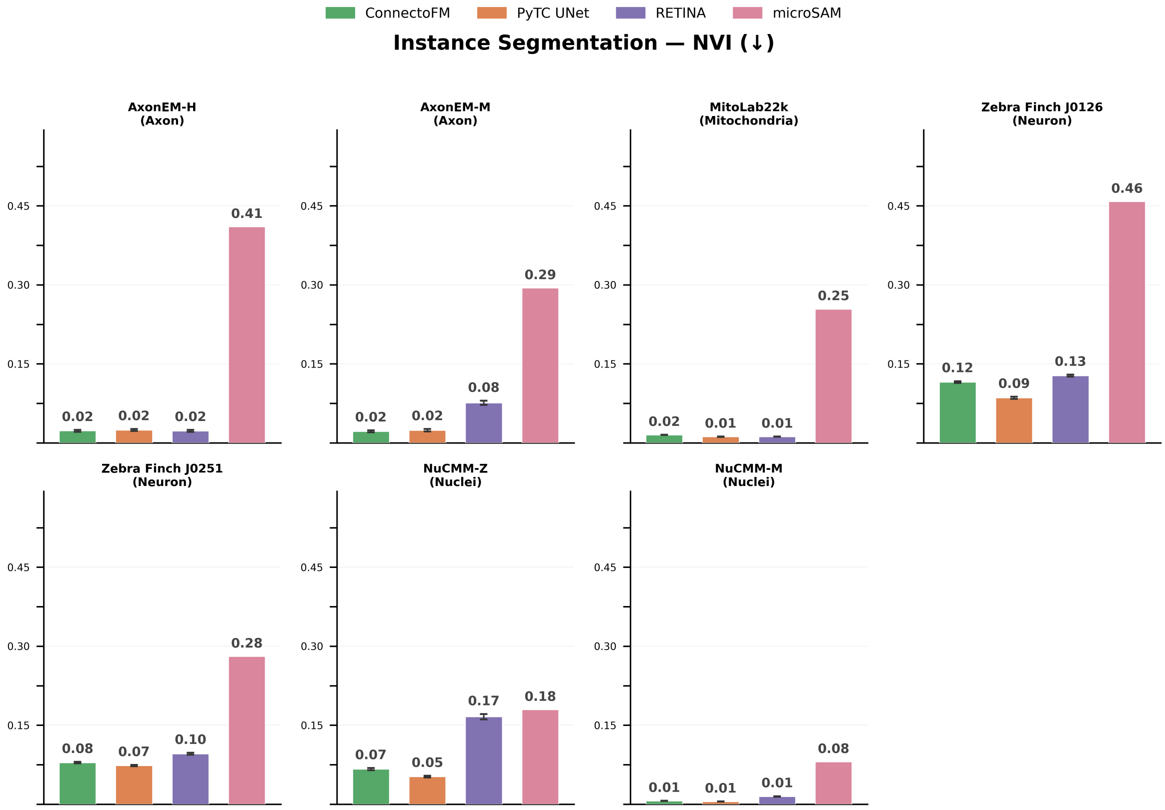


**Figure S8:** Comparison of NVI scores among ConnectoFM, PyTC UNet, RETINA, and microSAM in instance segmentation.

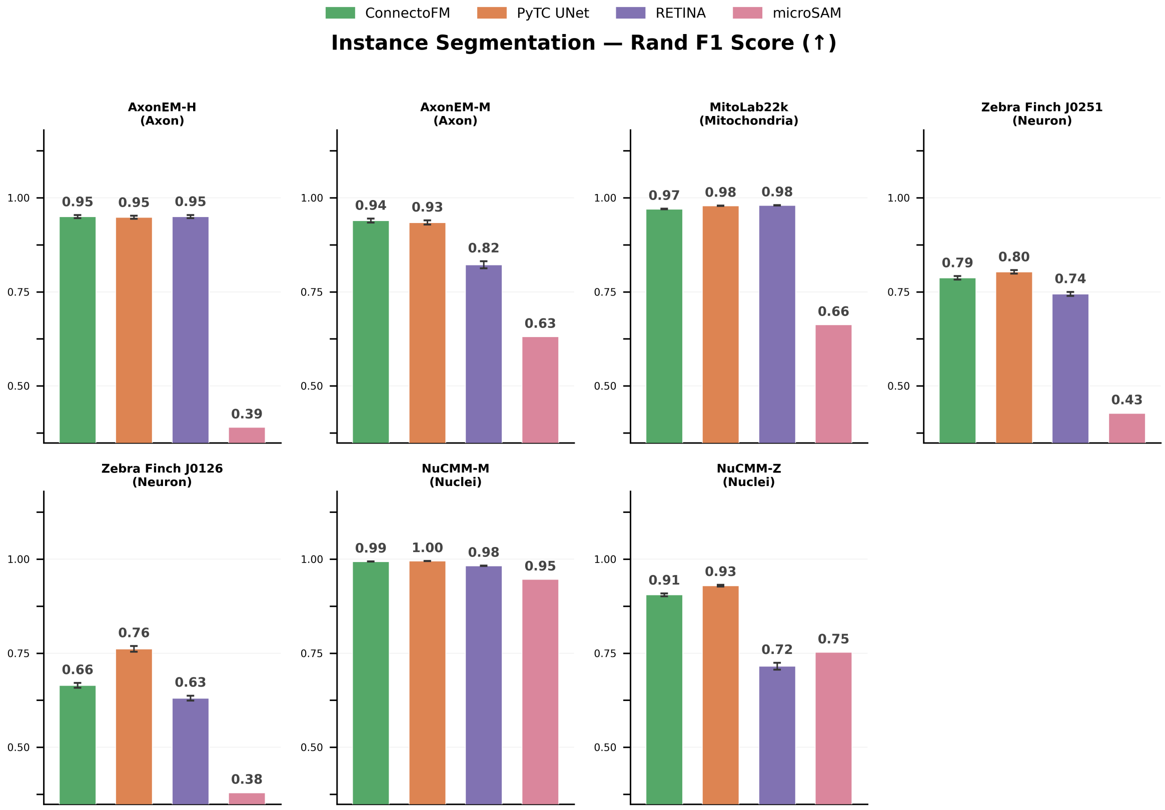


**Figure S9:** Comparison of Rand F1 Scores among ConnectoFM, PyTC UNet, RETINA, and microSAM in instance segmentation.

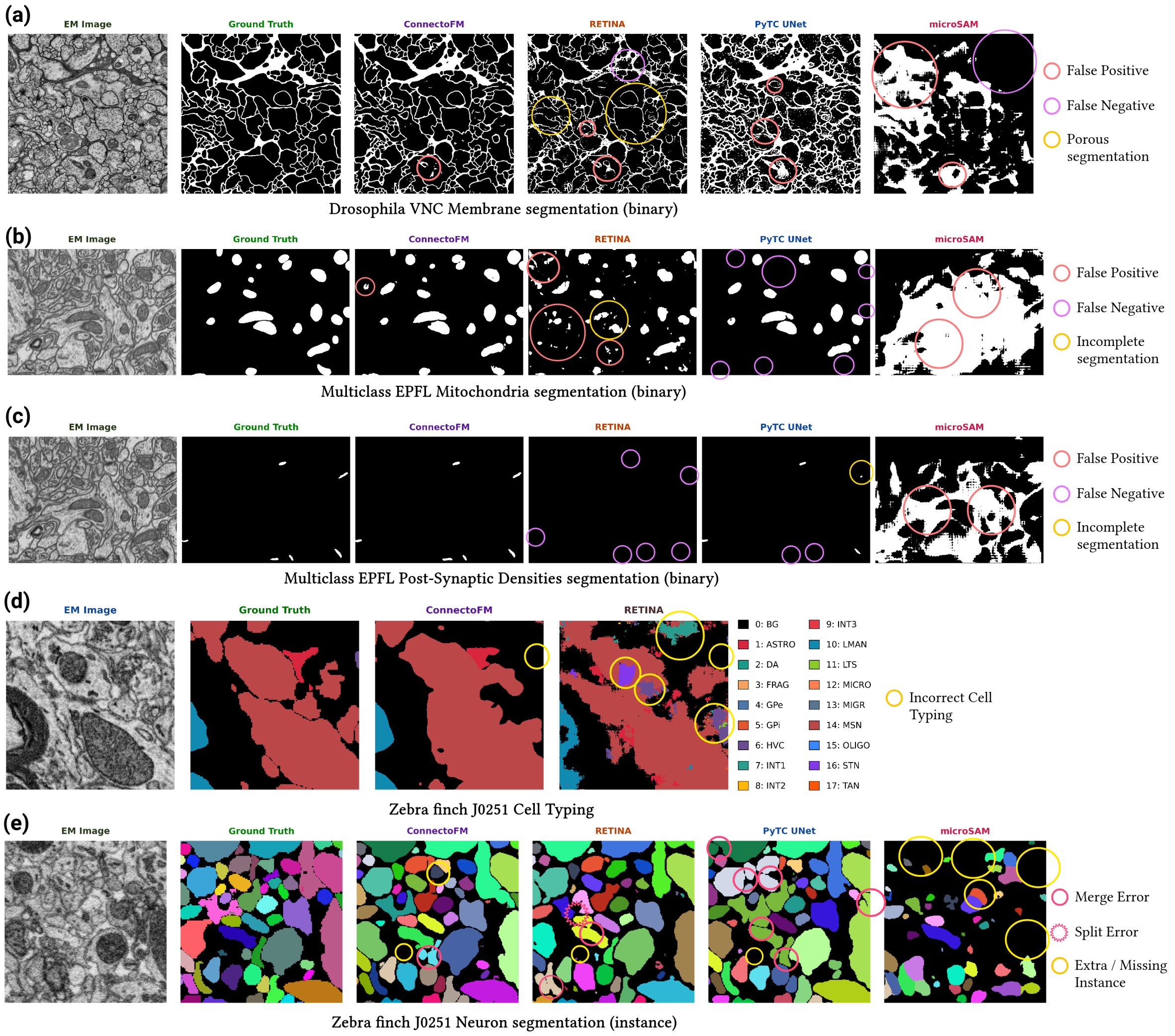


**Figure S10: Qualitative comparison of downstream performance between ConnectoFM and baseline methods. a–c,** Binary segmentation results for ConnectoFM, RETINA, PyTC UNet, and microSAM, with representative errors highlighted. ConnectoFM yields masks that more closely match the ground truth, whereas baseline methods show more frequent false positives, false negatives, and incomplete or porous segmentations. **a,** Membrane segmentation on the Drosophila VNC dataset. **b,** Mitochondria segmentation on the Multiclass EPFL dataset. **c,** Post-synaptic dendrite segmentation on the Multiclass EPFL dataset. **d,** Multiclass cell-typing predictions for ConnectoFM and RETINA on the Zebra Finch J0251 dataset, with representative incorrect cell-type assignments highlighted. **e,** Instance segmentation results for ConnectoFM, RETINA, PyTC UNet, and microSAM on the Zebra Finch J0251 dataset, with representative merge errors, split errors, and extra or missing instances indicated. This figure was created with BioRender.com.

#### References
